# *Piriformospora indica* employs host’s putrescine for growth promotion in plants

**DOI:** 10.1101/2021.01.19.427242

**Authors:** Anish Kundu, Abhimanyu Jogawat, Shruti Mishra, Pritha Kundu, Jyothilakshmi Vadassery

## Abstract

Growth promotion by endosymbiont *Piriformospora indica* has been observed in various plants; however, specific functional metabolites involved in *P. indica* mediated growth promotion are unknown. A GC-MS based untargeted metabolite analysis was used to identify *Solanum lycopersicum* metabolites altered during *P. indica* mediated growth promotion. Metabolomic analysis showed primary metabolites altered and specifically putrescine to be maximally induced in roots during the interaction. *P. indica* induced putrescine biosynthetic gene *SlADC1* in *S. lycopersicum* and acts via arginine decarboxylase (*ADC*) mediated pathway. *P. indica* did not promote growth in *Sladc*-VIGS (virus induced gene silencing of *SlADC* gene) lines of *S. lycopersicum* and when the ADC enzyme was inhibited with an inhibitor, DL-α-(Difluoromethyl) arginine. In Arabidopsis *adc* knock-out mutants, *P. indica* do not promote growth and this response was rescued upon exogenous application of putrescine. Putrescine promoted growth by elevation of auxin (indole-3-acetic acid) and gibberellin (GA_4_, GA_7_) levels in *S. lycopersicum*. Putrescine is also important for *P. indica* hyphal growth indicating that it is co-adapted by both host and microbe. Hence, we conclude that putrescine is an essential metabolite and its biosynthesis in plants is crucial for *P. indica* mediated growth promotion and fungal growth.

## Introduction

*Piriformospora indica* (syn. *Serendipita indica*, Basidiomycota) is a root endophytic fungus with a broad host range including monocots, dicots and eudicots (Varma et al., 1999; Johnson et al., 2018; Qiang et al., 2012). *Piriformospora indica* colonizes the root rhizodermis and cortex of many host plants including Arabidopsis, Maize, Tobacco, Barley, Rice and Poplar (Varma et al., 1999, Waller et al., 2005; Vadassery et al., 2008; Yadav et al., 2010; Jogawat et al., 2016). Increased nutrient uptake in the host plant is a major cause for *P. indica* induced plant growth promotion (Yadav et al., 2010; Bakshi et al., 2017; Rani et al., 2016; Prasad et al., 2019). *P. indica* also imparts biotic and abiotic stress tolerance by activating induced systemic resistance in shoots (Waller et al.,2005; Baltruschat et al., 2008; Sun et al.,2010; Jogawat et al., 2016). *P. indica* manipulates multiple plant hormone pathways during colonization: phytohormone, jasmonate and secondary metabolite, glucosinolate during early stages of interaction (Lahrmann et al., 2015), auxin and cytokinin during plant growth promotion in diverse plants (Xu *et al*., 2018). In many plants like barley, where *P. indica* causes cell death-associated colonization, the endophyte recruits GA signaling to degrade DELLAs and establish cell apoptosis susceptibility (Schäfer et al.,2009; Jacobs et al., 2011). Induced systemic resistance by *P.indica* in host plants is mediated by jasmonic acid signaling, GA signaling and cytoplasmic function of NPR1 (NONEXPRESSOR OF PATHOGENESIS-RELATED GENES 1) (Stein et al., 2008, Cosme et al., 2016). Plants also regulate the colonization through activation of basal defense pathway via cyclic nucleotide gated channel (CNGC19), which ensures controlled *P. indica* colonization (Jacobs et al., 2011; Jogawat et al., 2020). Elevated levels of plant-secondary metabolite, indole glucosinolate also restrict the propagation of *P.indica* and balance its growth on plant roots (Lahrmann et al., 2015). Global transcriptome and metabolome analysis has revealed the beneficial effects of *P. indica* on host plants such as Arabidopsis (Vahabi et al., 2015; Strehmel et al., 2016), barley (Molitor et al., 2011; Zuccaro et al., 2011; Ghabooli et al., 2013) and chinese cabbage (Hua et al., 2017). *P. indica*-mediated reprogramming of host plant’s transcriptome, proteome and metabolome under salt, water and drought stress has also been explored (Waller et al., 2005; Molitor et al., 2011; Alikhani et al., 2013; Ghabooli et al., 2013). In Arabidopsis, it has been observed that *P. indica* association primarily affects primary root metabolism and secondary metabolites like glucosinolates, oligolignols, and flavonoids (Strehmel *et al*., 2016). In chinese cabbage, *P. indica* alters γ-amino butyrate (GABA), oxylipin-family compounds, poly-saturated fatty acids, and auxin and its intermediates (Hua et al., 2017). No study so far assigns a functional role for a specific metabolite in *P. indica* mediated growth promotion across plants. Tomato (*Solanum lycopersicum* L.) is one of the most important vegetables grown worldwide with 177 million ton production (Saeed et al., 2019). However, Tomato has been least explored for its interaction with *P. indica* and beneficial effects. *P. indica* reduces the disease symptoms caused by the fungal pathogen *Verticillium dahliae* and repress the amount of Pepino mosaic virus in Tomato. It also increases Tomato fruit biomass in hydroponic culture and dry matter content (up to 20%) (Fakhro et al., 2010; Sarma et al., 2011). *P. indica* enhances the growth, Na^+^/K^+^ homeostasis, antioxidant enzymes and yield of tomato plants under normal and salt stress conditions (Abdelaziz et al., 2019). The metabolites involved in Tomato - *P. indica* interaction is poorly investigated despite the economic importance of this Solanaceous plant and growth enhancing role of *P. indica*. In this study, we investigated the metabolome alterations in *S. lycopersicum* during *P. indica* colonization to identify specific metabolites involved in *P. indica* mediated growth promotion.

Polyamines (PA) are low molecular weight carbon and nitrogen rich aliphatic compounds containing two or more amino groups that are essential for cell proliferation (Chen et al., 2019). In plants, PAs are mainly present in their free form as Putrescine (Put), Spermidine (Spd), and Spermine (Spm), soluble conjugated (to small molecules including phenolics) and insoluble bound form (bound to DNA, RNA, proteins) (Chen et al., 2019). Spermidine and spermine are synthesized from putrescine by sequential additions of amino propyl groups derived from decarboxylated S-adenosyl-Met (SAM) (Vuosku et al., 2012). Putrescine is synthesized from the amino acid ornithine and arginine by ADC (arginine decarboxylase) and ODC (ornithine decarboxylase) mediated pathways (Kusano et al., 2008; Liu et al., 2015). Polyamines including putrescine and spermidine are known to be involved in plant growth and development (Kusano et al., 2008, Takahasi & Kakehi, 2010, Liu et al., 2015), interactions of plants with growth promoting rhizobacteria (Valette et al., 2019) and also in plant defense against abiotic stresses (Kumria & Rajam, 2002; Cuevas et al., 2008; Alcázar et al., 2010). In this work, we dissect the metabolomic alterations induced by *P. indica* in *S. lycopersicum* roots and characterize the functional role of this highly induced metabolite.

## Results

### Endophytic fungus, *Piriformospora indica* stimulates root and shoot growth of *S. lycopersicum*

To study the time course of *P. indica* growth and colonization pattern in tomato, we conducted a growth promotion assay at different days post inoculation (dpi). Chlamydospores were visible in the root cortex at 10 dpi with *P. indica* (Fig. S1), though no growth promotion was observed. After 30 dpi, growth promotion by *P. indica* was first observed (Fig. S2A) and at 40 dpi, the inoculated plants showed maximum growth promotion (Fig. 1A). For tracking of colonization, a green fluorescent protein (GFP)-tagged *P. indica* strain was utilized (Hilbert et al., 2012, Jogawat et al., 2020), and we observed both the chlamydospores and fungal hyphae in root cortex at 40 dpi, indicating endophytic colonization (Fig. 1B). The first significant stimulation of growth parameters in *P. indica*-treated plants were observed after 30 dpi and maximum at 40 dpi as compared to the control plants (Fig. 1C). At this stage, root fresh weight (Fig.1D, 1E), shoot and root length (Fig. S2B, S2C) were also found to be significantly increased in *P. indica* treated plants. We quantified the fungal DNA content in the roots and observed around 8 fold increase at 30 dpi and around 30 fold increase at 40 dpi, over the control (0 dpi is the first day of fungal inoculation) (Fig. 1F). As at 40 dpi maximum fungal colonization and growth promotion was observed, we selected this time point for further investigations.

**Figure 1.**
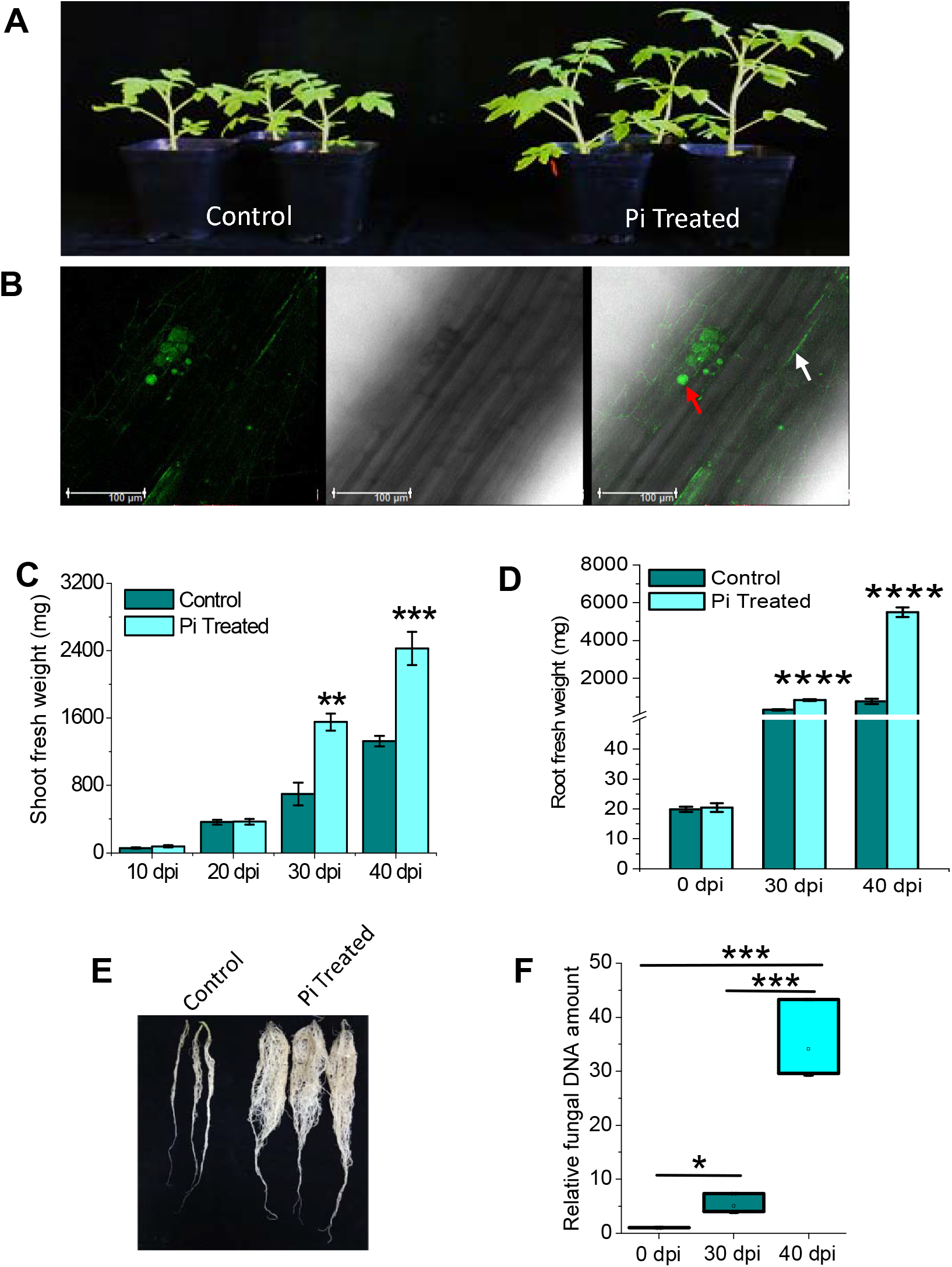
Effect of *P. indica* (Pi) treatment on phenotype of tomato (*S. lycopersicum*) plants in a time course (A) Representative *S. lycopersicum* shoot growth in non-inoculated (left tray) and *P. indica* inoculated (right tray) pots. The experiment was conducted for three independent times. (B) GFP-labeled *P. indica* colonization pattern after 40 dpi in tomato root; (left) fluorescence, (middle) bright field, (right) merged image. Red and white arrows indicate *P. indica* chlamydospores and mycelia respectively. (C) Mean of shoot fresh weight ± S.E. (n = 10) at different time points after upon *P. indica* inoculation. (D) Quantification of root growth upon 30 dpi and 40 dpi of *P. indica* inoculation (n = 10). (E) Visualization of root growth upon 40 days of *P. indica* inoculation. The figure is the best representative of three independent experimental sets. (F) Quantification of *P. indica* colonization (n = 4) in roots after 30 dpi and 40 dpi of inoculation. 0 dpi denotes first day of inoculation. Relative fungal colonization was measured by subtracting the C_T_ values of *P. indica Tef1* from C_T_ values of tomato *UBI3* gene. Significance analysis was done by unpaired *t*-test; \**P*<0.05, \*\**P*<0.01, \*\*\**P*<0.001, \*\*\*\**P*<0.0001

### *Piriformospora indica* colonization alters leaf and root metabolites of *S. lycopersicum*

To explore the global metabolome alteration in tomato upon *P. indica* colonization, we conducted untargeted gas chromatography-mass spectrometry (GC-MS) analysis at 40 dpi in both shoots and roots separately. We also profiled the *P. indica* hyphal (mycelia) metabolome, to distinguish tomato-specific metabolites. Metabolite profiling revealed in total 425 mass signals (124 for leaf, 163 for root and 138 for *P. indica* mycelia), among which 55 were identified and annotated in leaf, 70 in root and 102 in *P. indica* mycelia (Fig. 2A, Table S2). A comparative analysis of annotated metabolites revealed 76 mycelia specific, 22 root specific and 9 leaf specific metabolites (Fig. 2B). Root and leaf shared 45 metabolites, root and mycelia shared 3 metabolites, leaf and mycelia shared only 1 metabolite while 22 metabolites are shared by root, leaf and mycelia (Fig. 2B, Table S3). The annotated metabolites in tomato covered a broad range of primary metabolites including sugars and amino acids, while only three secondary metabolites (caffeic acid, chlorogenic acid and benzoic acid) were identified (Table S2). Normalized peaks of annotated leaf and root metabolites were used for ‘Pearson correlation coefficient’ (PCC) calculation, which revealed a strong positive correlation between control and treated data sets of leaf (PCC = 0.89) and root (PCC = 0.87) metabolites (Fig. S3a and S3b). To get a global view of fold changes of metabolites shared by leaf and root we created a Pearson’s correlation based clustered heat map (Fig. 2C). Metabolite fold changes in leaf and root showed negatively correlated clusters. We further analyzed correlation among leaf and root on the basis on metabolite fold changes and constructed a correlation network where leaf and root showed negative correlation (Fig 2D). This indicates *P. indica* infestation in root alters root and leaf metabolome reversibly.

**Figure 2.**
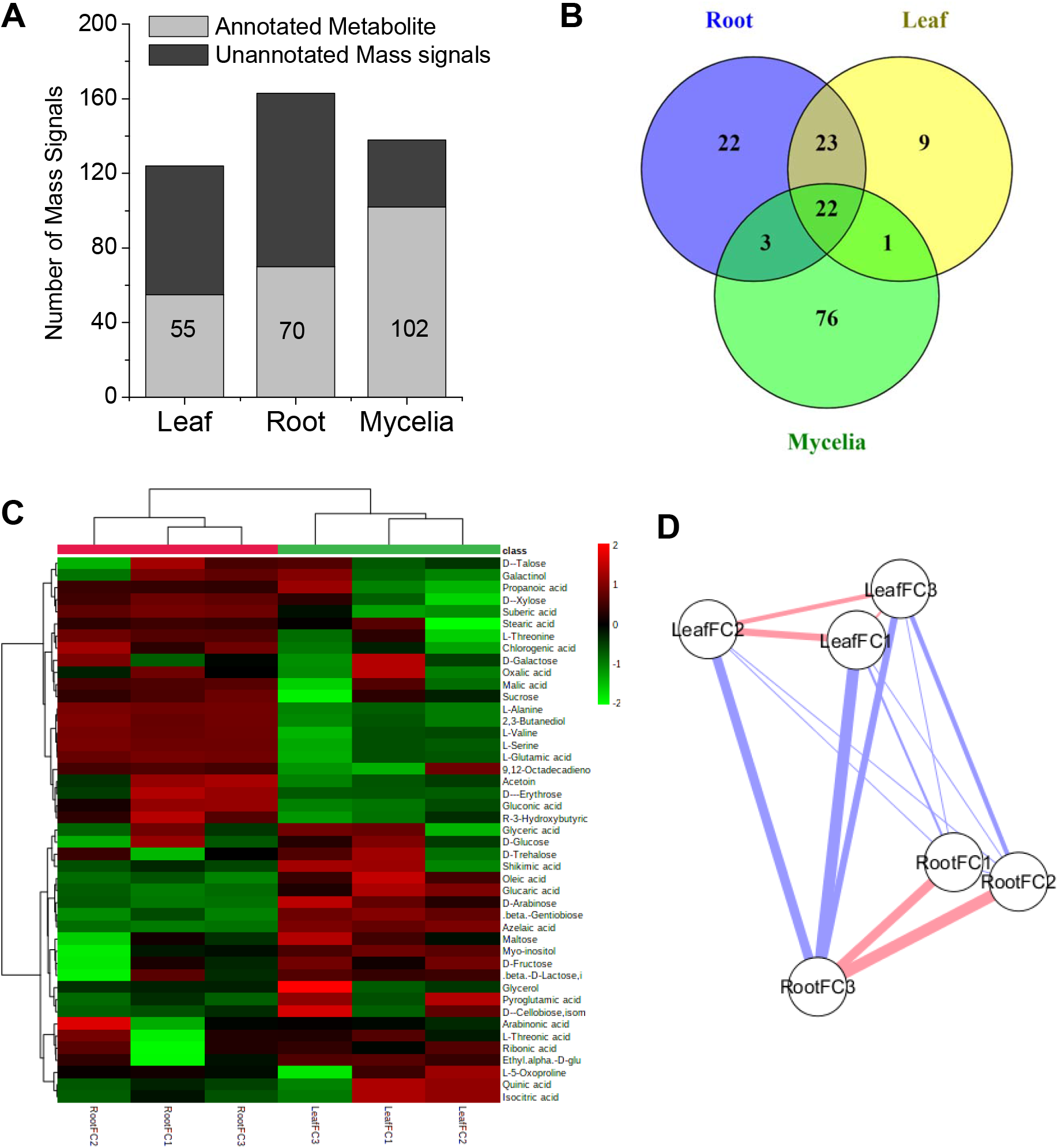
Alteration of annotated metabolites in tomato leaf, root and *P. indica* mycelia. (A) Mean of total number of GC-MS mass signals (n = 3, each replicate is the pool of three individual plants) compared to number of identified and annotated metabolites detected in leaf and root (control and *P. indica* treated) of tomato and *P. indica* mycelia. (B) Venn diagram to show comparative metabolite profile and number of specific and common metabolites detected in leaf, root and *P. indica* mycelia. (C) Heat map with Pearson’s correlation based clustering (algorithm: complete) of Log_2_ fold change of 45 common metabolites in *S. lycopersicum* root and leaf. Scale shows change values. Heat map was generated in MetaboAnalyst 4.0. (D) Pearson correlation network in between root and leaf (n = 3) on the basis of Log_2_ fold change values of the 45 common metabolites in leaf and roots. Nodes represent leaves and roots, edges represent correlations (blue: negative correlation; red positive correlation); thickness of the edges represents correlation strength. Pearson correlations was calculated in MetaboAnalyst correlation algorithm and the network is generated in Cytoscape 3.2.1 aided with MetScape.

### Putrescine is a central metabolite altered in tomato roots upon *P. indica* colonization

Metabolite fold changes (fold of up- or down-regulation of each metabolite’s normalized peak area in *P. indica* colonized plant compared to control plant) were calculated from GC-MS data-set for root and leaf specific metabolites (Table S4). In leaf, volcano plot showed up-regulation of five metabolites i.e. 12-hydroxyoctadecanoic acid, glucaric acid, β-D-lactose, arabinose, fructose and down-regulation of four metabolites i.e. L-alanine, chlorogenic acid, azelaic acid and 2, 3-butanediol (Fig. S4). Metabolites having an alteration cut-off value of *P*<0.05 and FC>1.5 were considered as ‘significantly altered’ in the volcano plot.

As *P. indica* is a root infecting endophyte we decided to specifically look at root specific metabolites. First we performed a partial least squares-discriminant analysis (PLS-DA) with all root metabolite data. Control and *P. indica* infested roots were classified in two different groups indicating a clear divergence in their metabolite levels (Fig. 3A). A variables of importance (VIP)-score plot was generated from the PLS-DA model, which showed that metabolite putrescine has highest VIP score (>1.38) indicating its significance among altered metabolites (Fig. 3B). Volcano plot showed up-regulation of eight metabolites i.e. Putrescine (1’), Gluconic acid (2’), Glucaric acid (3’), L-Alanine (4’), Propanoic acid (5’), L-Glutamic acid (6’), Lactic acid (7’) and Acetoin (8’) and also significant down-regulation of eight metabolites i.e. Benzoic acid (9’), Myo-inositol (10’), Azelaic acid (11’), Phenylpyruvic acid (12’), Fumaric acid (13’), L-Valine (14’), 9,12-Octadecadienoic acid (15’) and Shikimic acid (16’) (Fig. 3C and Fig. S5A and B). In roots, putrescine showed maximum increase and benzoic acid showed maximum decrease (Fig. 3C and Fig. S5A and B). In GC-MS analysis, putrescine was not detected in *P. indica* mycelia, therefore, for confirmation we performed LC-MS/MS analysis, where putrescine was observed in very low intensity (Fig 3D), which indicates its increase is majorly root specific. We also analyzed the root metabolite data with metabolite marker selection approach (Zang et al., 2013) and performed orthogonal partial least-squares to latent structures discriminant analysis (OPLS-DA) and generated an S-plot. S-plot showed putrescine to have highest reliability and magnitude (p[1]6.1; p(corr)[1] 0.997) to be a metabolite marker (Fig. S5C). Therefore, putrescine has been considered as the most significant metabolite during interaction in between *P. indica* and *S. lycopersicum* root.

**Figure 3.**
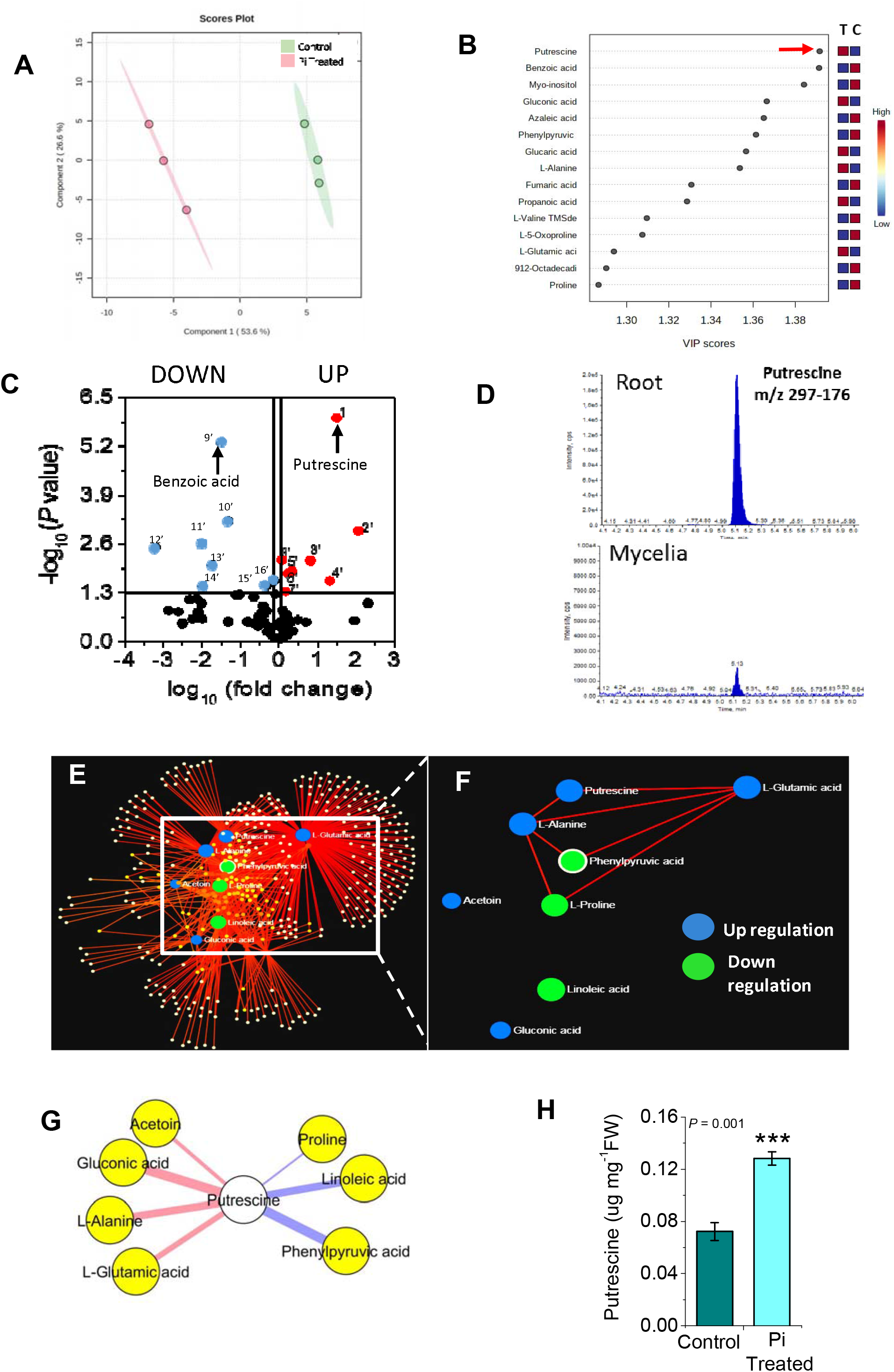
Analysis of differential alteration of root metabolite (A) PLS-DA score plot of control and *P. indica* treated root samples on the basis of 77 metabolite’s normalized peak areas. (B) VIP (variables of importance) score plot shows top fifteen variables (metabolites) of importance in the root. Red arrow indicates putrescine with highest VIP score. (C) Volcano plot shows up-regulation (Red) of eight metabolites and down regulation (Blue) of eight metabolites in root. Point denoted as 1’ shows maximum up-regulated putrescine and 9’ shows maximum down regulated benzoic acid. Numbering of the denoted metabolites are described in figure S5.(D) Comparative LC-MS/MS XIC showing putrescine (Q1-Q3: m/z 297-176) peaks in root and mycelia. (E) Metabolite-metabolite interaction network constructed with the fold change values of significantly altered root metabolites, which predicted 404 plausible nodes (metabolites) and 830 plausible edges (interactions). (F) Zoomed view of interaction network in between up-regulated and down-regulated metabolites. (G) Correlation network of interacting upregulated and down regulated metabolites with putrescine. (H) Mean ± SE of absolute amount of putrescine in control and *P. indica* colonized *S. lycopersicum* roots after 40 dpi. Significance analysis was done by unpaired *t*-test; \**P*<0.05, \*\**P*<0.01, \*\*\**P*<0.005

Pathway analysis with significantly altered (*P*<0.05) annotated root metabolite data set showed a plausible involvement of 46 metabolic processes (Fig. S6 and Table S5). We considered pathways having *P*<0.05, impact>0.1 as truly impacted pathways. Among the top three pathways alanine, aspartate and glutamate metabolism showed highest impact (*P* = 2.48e^−07^, impact = 0.327), followed by tyrosine metabolism (P = 5.92e^−06^, impact = 0.108) and arginine and proline metabolism (*P* = 2.17e^−05^, impact = 0.144). Metabolite-metabolite interaction network predicted 404 nodes (metabolites) and 830 edges (interactions) (Fig. 3E and Table S6). The up-regulated putrescine, L-glutamic acid and L-alanine have direct interaction with each other and with down-regulated phenylpyruvic acid and L-proline which indicates coordination among them. But no interaction was found with the other two up-regulated metabolites (acetoin and gluconic acid) and down-regulated, octadecadienoic acid (Fig. 3F). These results further signify the importance of highly upregulated putrescine to be majorly involved in metabolic interaction with other up-regulated molecules during tomato and *P. indica* interaction. For confirmation of these interactions, we performed a Pearson’s correlation based network analysis in between the interacting metabolites, and observed putrescine is positively correlated in alteration with the acetoin, gluconic acid, L-alanine and L-glutamic acid whereas, negatively correlated with proline, linoleic acid and phenylpyruvic acid (Fig 3G). Interestingly, gluconic acid showed strong positive correlation (PCC = 0.70583) but without significance (*P* = 0.117). Finally, we did an absolute quantification of putrescine independently using LC-MS/MS method, to show a highly significant elevation of putrescine content in *S. lycopersicum* roots upon *P. indica* colonization (Fig. 3H).

### *P. indica* induced putrescine biosynthetic gene in *S. lycopersicum*

As putrescine level significantly increased in *S. lycopersicum* roots upon *P. indica* colonization, we examined the expression levels of putrescine biosynthetic genes. Putrescine is synthesized from the amino acids arginine and ornithine by ADC and ODC mediated pathways and genes involved in the process are arginine decarboxylase (*SlADC1 and SlADC2*) and ornithine decarboxylase (*SlODC1, SlODC2 and SlODC3*) (Liu et al., 2018). We checked all five gene (*SlADC* and *SlODC*) transcript levels in *S. lycopersicum* colonized by *P. indica*. We found significantly increased transcript level only for *SlADC1*, and no transcript induction was observed for any *SlODC* (Fig. 4A), which confirmed that *P. indica* induced putrescine biosynthesis is through arginine decarboxylase (*ADC*) mediated pathway.

**Figure 4.**
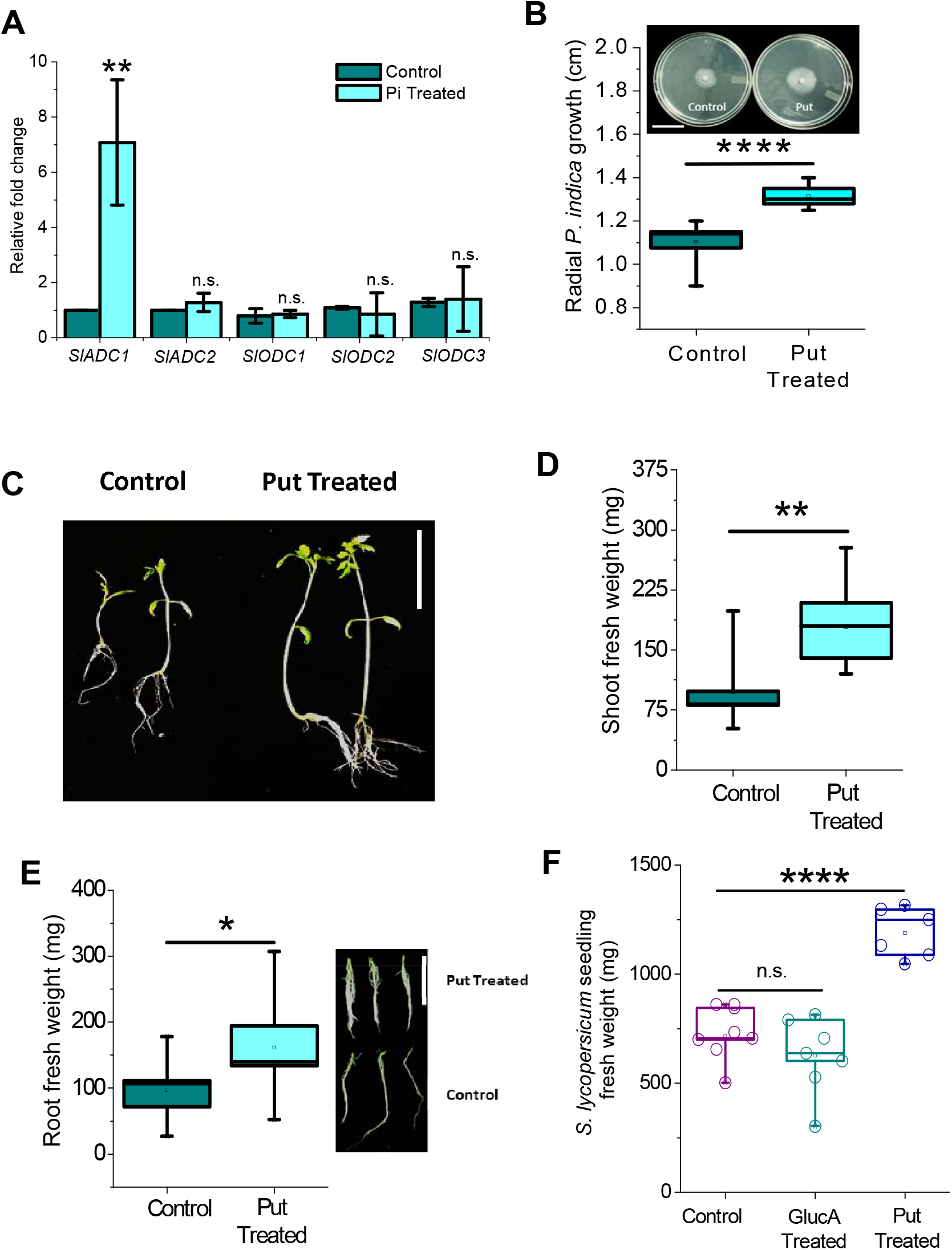
Effect of exogenous putrescine on *P. indica* and *S. lycopersicum* growth and alteration of putrescine biosynthetic genes in *S. lycopersicum* upon *P. indica (Pi*) colonization. (A) Mean ± SE of arginine decarboxylase (*SlADC*) and ornithine decarboxylase (*SlODC*) transcript levels after 40 dpi of *P. indica* inoculation (n = 4). Controls of *SlADC1* and *SlADC2* has error bars merged with the bar’s out line as they have small values. (B) Quantification of radial growth of *P. indica* upon 10 μM putrescine treatment (n = 10). Inset shows the visualization of the radial growth. (C) Visualization of phenotype of growth in *S. lycopersicum* seedlings upon 10 μM putrescine treatment. Scale bar: ~4.5 cm. The figure is the representative of 10 replicates. Quantification of (D) shoot fresh weight and (E) root fresh weight of *S. lycopersicum* and visualization of root growth upon 10 μM putrescine treatment (n = 10). Scale bar: ~4 cm. (F) Fresh weight of *S. lycopersicum* seedlings upon 10 μM Gluconic Acid (GlucA) and 10 μM Putrescine (Put) Treatment. Significance analysis was done by unpaired *t*-test; \**P*<0.05, \*\**P*<0.01, \*\*\**P*<0.001, \*\*\*\**P*<0.0001.

### Putrescine induced growth of both *P. indica* and *S. lycopersicum*

Polyamines were previously reported to have role in fungal cell differentiation and development (Stevens and Winther, 1979, Ruiz-Herrera, 1993). We checked whether putrescine, has a role in *P. indica* growth *per se*, thus providing an advantage for its up-regulation. Five different putrescine concentrations (5 μM-100 μM) were tested to check its effect on the radial growth of *P. indica*. It was observed that 10 μM putrescine significantly induced *P. indica* radial growth, while putrescine concentration higher than 10 μM did not induce further radial growth (Fig. 4C; Fig. S8A). For additional confirmation, we applied 10 μM putrescine in the liquid *P. indica* culture, which showed significant induction in both the fresh and dry biomass (Fig. S8 B, C) after 14 days of treatment. These results suggest putrescine enhances growth of *P. indica*. We attempted to identify the specific role of putrescine in *S. lycopersicum* growth, and therefore, exogenously treated 5 day old tomato seedlings. After 21 days of treatment, we found significant induction in multiple growth parameters of *S. lycopersicum* (Fig. 4D), including shoot fresh weight (Fig. 4E), root fresh weight (Fig. 4F) and shoot length (Fig. S7). Hence, putrescine is involved in growth promotion of *S. lycopersicum*. In the volcano plot, apart from putrescine, gluconic acid is also increased significantly and was second most significantly upregulated metabolite. Therefore, we further examined the comparative effect of putrescine and gluconic acid in 5 day old *S. lycopersicum* seedlings. 21 days of treatment, 10 μM gluconic acid did not show any growth promotion, whereas 10 μM putrescine promoted the growth significantly (Fig. 4A).

### Putrescine induced growth promotion in *S. lycopersicum* by elevating auxin and gibberellin biosynthesis

Auxin and cytokinin are major growth hormones involved in plant growth promotion response of *P. indica* in diverse plants (Sirrenberg et al., 2007; Vadassery et al., 2008; Meents et al., 2019). *P. indica* needs also induces the GA biosynthesis during colonization (Cosme et al., 2016). Hence we addressed the possible involvement of putrescine to alter these growth hormone levels in *S. lycopersicum*. We measured indole-3-acetic acid (IAA), five gibberellins (GA_1_, GA_3_, GA_4_, GA_7_ and GA_8_) and nine cytokinins (tZ, *trans*-zeatin; tZR, *trans*-zeatin riboside; DHZR, dihydrozeatin riboside; tZROG, *trans*-zeatin riboside-O-glucoside; tZ7G, *trans*-zeatin-7-glucoside; DHZ, dihydrozeatin; DHZROG, dihydrozeatin riboside-*O*-glucoside; DHZOG, dihydrozeatin-*O*-glucoside; iP, isopentenyladenine) in the putrescine treated *S. lycopersicum* seedlings. It was observed that IAA level increased significantly in the putrescine-treated seedlings (Fig. 5A). Similarly, GA_4_ and GA_7_ were also significantly increased (Fig. 5B). No rise were observed in the cytokinin levels, instead DHZR, tZ7G and DHZ levels decreased significantly (Fig. 5C). This result signifies the involvement of IAA and GAs in putrescine induced growth promotion.

**Figure 5.**
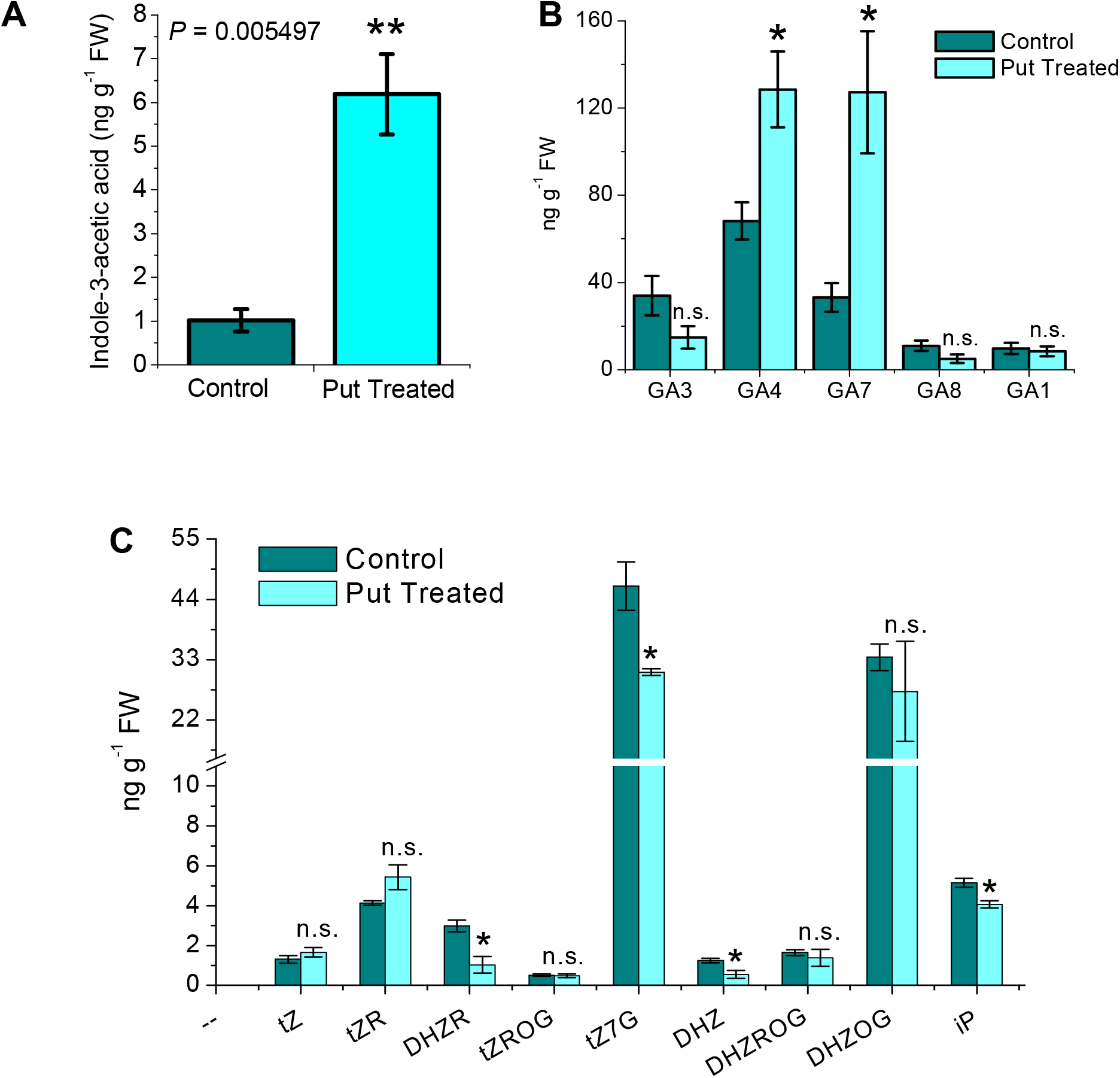
Effect of exogenous putrescine on growth phytohormone levels in *S. lycopersicum*. (A) Mean ± SE (n = 4) of indole-3-acetic acid (IAA) (B) Mean ± SE (n = 4) of gibberellins (GA_1_, GA_3_, GA_4_, GA_7_ and GA_8_) and (C) Mean ± SE (n = 4) of different cytokinins in control and 10 μM putrescine treated *S. lycopersicum* seedlings. tZ, *trans*-zeatin; tZR, *trans*-zeatin riboside; DHZR, dihydrozeatin riboside; tZROG, *trans*-zeatin riboside-*O*-glucoside; tZ7G, *trans*-zeatin-7-glucoside; DHZ, dihydrozeatin; DHZROG, dihydrozeatin riboside-*O*-glucoside; DHZOG, dihydrozeatin-*O*-glucoside; iP, isopentenyladenine.

### Putrescine biosynthesis is crucial for *P. indica* mediated growth promotion in *S. lycopersicum* and *A. thaliana*

We then tested if putrescine biosynthesis functionally plays a role in *P. indica-mediated* growth induction. A *S. lycopersicum* line was generated where *SlADC1* was silenced by virus-induced gene silencing (VIGS) approach (Fig. S9 and S10). The line was confirmed by checking *SlADC1* transcript level compared to the control i.e. empty vector transformed (EV) plants. After 7 days and 14 days post agro infiltration of VIGS construct, they were silenced upto 97.41% and 85.93% respectively (Fig. 6A). At this stage putrescine content was measured and found to be significantly reduced (Fig. 6B), which confirmed the *ADC* silencing. Both EV and *adc1*-VIGS plants under control conditions showed no growth differences, However, upon co-cultivation with *P. indica* for 40 dpi, both the shoots and roots of *adc1*-VIGS plants showed no growth promotion when compared to EV+ *P. indica* plants (Figs. 6C, 6D and 6E). In order to reconfirm the importance of *ADC*-mediated putrescine biosynthesis, we treated the *S. lycopersicum* seedlings in the media with DL-α-(Difluoromethyl) arginine (DFMA), an irreversible inhibitor of ADC enzyme, which was previously reported to efficiently reduce cellular putrescine level in *S. lycopersicum* (Fernández-Crespo et al., 2015). It was observed that *P. indica* showed reduced growth promotion in the DFMA treated *S. lycopersicum* seedlings compared to control. Both the shoot and root fresh weights were reduced in the DFMA treated seedlings (Fig. S11A and B). On the other hand, 2 weeks of DFMA treatment did not show any inhibitory activity on *P. indica* growth (Fig. S11C and D). These results reconfirm importance of ADC1 in *P. indica*-mediated growth promotion in *S. lycopersicum*.

**Figure 6.**
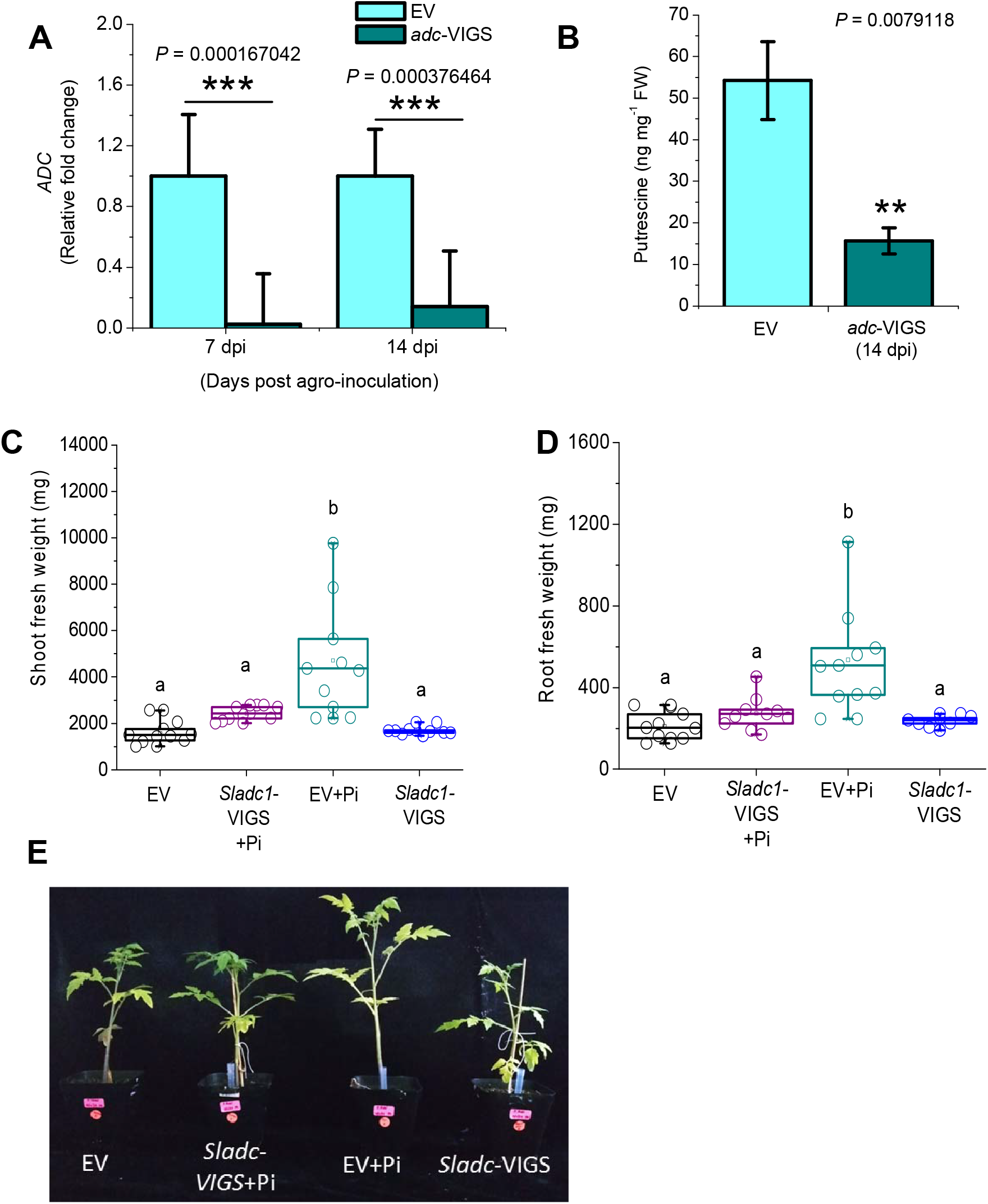
*P. indica* mediated growth in *S. lycopersicum adc1*-VIGS plants. (A) Mean + SE of arginine decarboxylase (ADC1) transcript levels in EV and *adc*-VIGS *S. lycopersicum* after 7 dpi and 14 dpi (n = 4). Silencing efficiency at 7 dpi is 97.41% and at 14 dpi is 85.93%. (B) Mean ± SE of putrescine content in EV and *Sladc1-VIGS* plants (n = 4). Significance analysis was done by unpaired *t*-test; \**P*<0.05, \*\**P*<0.01, \*\*\**P*<0.001. Quantification of the (C) root fresh weight (n = 10) and (D) shoot fresh weight (n = 8-10) in EV and *Sladc1*-VIGS. Significance analysis was done by analysis of variance (ANOVA) followed by Tukey’s test. Different letters denoted significant differences. (E) Visualization of phenotypic changes and growth promotion of EV and *adc1*-VIGS plants after 40 dpi of *P. indica* infestation. This figure is a representative of 8-10 biological replicates of each of the plants.

To study the role of putrescine biosynthesis across plants, we checked its effect on Arabidopsis. *A. thaliana* seedlings grown on media with 10 μM putrescine showed a significant induction of both fresh weight (*P*< 0.01) (Fig. 7A and B) and root length (*P*< 0.001) (Fig. S12A). Previously, it was reported that overproduction of putrescine and other polyamines reduced the growth in Arabidopsis (Alcázar et al., 2005), but here the results indicate that exogenous application of low concentration (10 μM) of putrescine stimulates growth in Arabidopsis. For functional characterization of the *adc* mutants in Arabidopsis we used the previously reported lines (Cuevas *et al*., 2018). *adc1-2* (SALK_085350C), *adc 1-2*(CS9658)*, adc1-3*(CS9657), *adc2-3*(CS9659) and *adc2-4*(CS9660) (Fig. S12B and C) and treated them with *P. indica*. It was previously demonstrated that *adc1-2, adc1-3, adc2-3* and *adc 2-4* accumulated significantly less amount of free putrescine (Cuevas *et al*., 2018). In our experiment, we found that after 14 dpi, fresh weight and root length were significantly induced only in wild type, whereas *adc* mutants did not show any growth promotion (Fig. 5C, D; Fig. S12D). This result signifies that *P. indica* fails to induce plant’s growth when putrescine biosynthesis is impaired. In a parallel experiment we performed a complementation assay, where *adc* mutants (*adc 1-2, adc 1-3,adc 2-3* and *adc 2-4*) were treated with *P. indica* in a media containing 10 μM putrescine. After 14 dpi of *P. indica* inoculation, both fresh weight (Fig. 7C and D) and root length (Fig. S12D) were increased in putrescine complemented mutants, which signifies that exogenous putrescine complemented the mutant phenotype. These result confirms that putrescine is required for *P. indica*-mediated growth induction in both Arabidopsis and *S. lycopersicum*.

**Figure 7.**
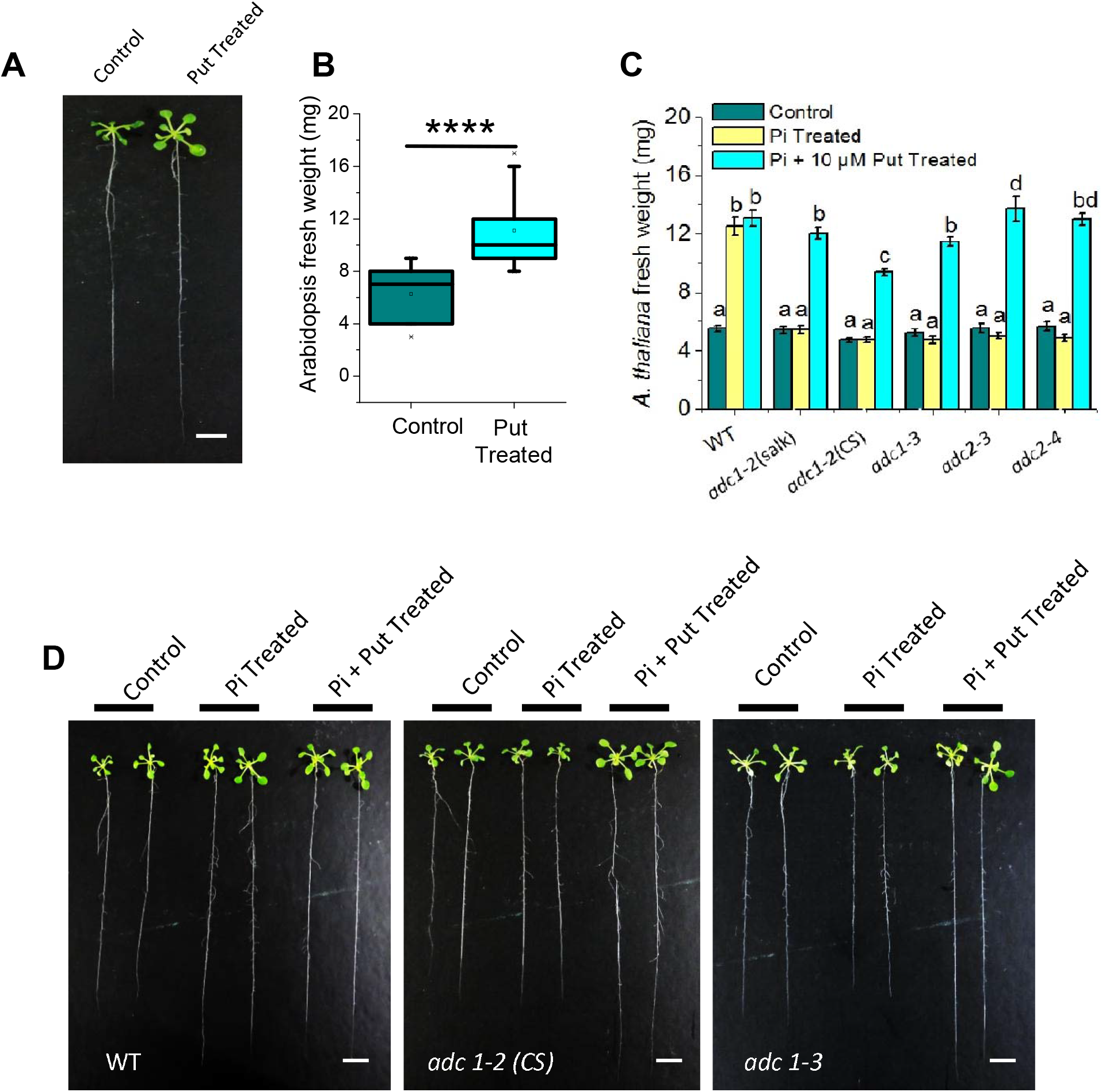
Effect of putrescine on Arabidopsis growth (A) Representative figure of 14 days old Arabidopsis seedlings treated with 10 μM putrescine (n=12) (B) Quantification of fresh weight of seven days old Arabidopsis seedlings upon 10μM putrescine treatment (n = 12). Significant analysis was done by unpaired t-test (****P<0.0001). (C) Growth promotion assay in putrescine biosynthetic mutants along with gain-of-function assay by putrescine supplementation. Mean ± SE (n = 22 - 30). Significance analysis was done by ANOVA (*P<0.05). (D) Representative figure of 22-30 biological replicates of putrescine induced *P. indica* mediated growth promotion assay in two *adc* mutant lines of Arabidopsis (14 days old). Scale bar: 0.4 cm. The seedlings were transferred from plates to a black surface for photograph

## Discussion

Mutualistic interactions of plants with *P. indica* can enhance growth through mechanisms such as nutrient uptake (phosphate and nitrate uptake, sugar transport), phytohormone production (auxin and cytokinin) and indirectly through induced systemic resistance (Yadav *et al.,2010*; Sherameti *et al*., 2005; Rani *et al*., 2016; Vadassery *et al*., 2009; Sirrenberg *et al*., 2007; Vahabi *et al*., 2018). Plant roots release many metabolites into the rhizosphere to kick-start symbiotic interactions, including flavonoids that act as chemo-attractants for rhizobial bacteria (Liu *et al*., 2016; Oldroyd 2013) and strigolactones for mycorrhizal fungi (Besserer *et al*., 2006). Roots also accumulate metabolites and transport it to shoots as in the case of blumenol C-glucoside accumulation upon mycorrhizal colonization in *Nicotiana attenuata* (Wang *et al*., 2018). Host plant metabolites that are responsible for *P. indica* mediated plant growth promotion are not known. In Chinese cabbage roots, *P. indica* stimulated the synthesis of metabolites involved in the tryptophan and phenylalanine metabolism and γ-aminobutyrate (GABA). Tryptophan and indole metabolism were speculated to be used for *de novo* biosynthesis of auxin in *P. indica*-colonized roots, facilitating growth promotion in Chinese cabbage. (Hua *et al*., 2017). Here, we report the identification and functional characterization of a specific metabolite, putrescine, to be involved in growth promotion response of *S. lycopersicum* upon *P. indica* colonization.

Metabolite analysis shows that host pathways in roots targeted by *P. indica* belong to primary metabolism, including amino acids and polyamines In tomato roots the polyamine, putrescine was the highest induced metabolite upon *P. indica* colonization after establishment of the symbiotic interaction at 40 dpi. Multivariate statistical analysis and metabolite marker finding approach also suggested putrescine as mostly highlighted candidate among the eight upregulated root metabolites. Out of these upregulated root metabolites, alteration of putrescine, L-glutamic acid, L-alanine, phenylpyruvic acid and L-proline directly interact with each other in the metabolite-metabolite interaction network analysis. Pathway analysis with significantly altered root metabolites also revealed amino acid, arginine and glutathione metabolism to be altered in *S. lycopersicum* root. Arginine and glutathione metabolisms regulate putrescine and glutamic acid level in the cell respectively; arginine is the precursor in putrescine biosynthesis, while glutamic acid is the precursor as well as degradation product of the glutathione (Mo *et al*., 2015; Liu *et al*., 2010). In previous reports it was also demonstrated that during antioxidant based defense, exogenous polyamines induces glutathione level to reduce the overproduction of reactive oxygen species (Nahar *et al*., 2016 a, b). Significant impact of arginine metabolism indicates the primary involvement of putrescine biosynthetic pathway during *P. indica* interaction with *S. lycopersicum*. In our study, transcript level of putrescine biosynthetic gene *SlADC1*, was induced in *P. indica* colonized root. The knock-down of *adc* genes in Arabidopsis and tomato also resulted in loss of growth promotion response. Hence *P. indica* induces putrescine biosynthesis only by arginine decarboxylase mediated pathway which is functionally important for growth promotion. In tomato, *ADC* and *ODC* have differential tissue expression and *ADC* is root expressed (Acosta *et al*., 2005; Kwak & Lee, 2001). *P.indica* also produces putrescine in a very less concentration in whole mycleia, however this does not complement the phenotype of *adc* mutants suggesting the critical role of plant induced putrescine. However, our study does not rule out involvement of additional functional metabolites at different stages of *P. indica* colonization.

Polyamines are also known to be involved in plant embryogenesis and growth (Kusano *et al*., 2008, Takahasi & Kakehi, 2010, Liu *et al*., 2015). Though, overproduction of putrescine and other polyamines reduced the growth in Arabidopsis (Alcázar et al., 2005), its affirmative effect on root cultures of chicory is reported (Bais *et al*., 2001). Thermospermin plays a role in stem elongation of Arabidopsis (Knott et al., 2007). A double mutant of putrescine biosynthetic gene, arginine decarboxylase (*adc1^−^/adc2^−^*) and spermidine biosynthetic gene, spermidine synthase (*spds1^−^/spds2^−^*) showed lethal defect in embryo development in Arabidopsis (Imai *et al*., 2004; Urano *et al*., 2004). Polyamines, including putrescine, are known to be induced upon abiotic and biotic stress (Liu *et al*., 2015; Minocha *et al*., 2014; Seifi & Shelp 2019). Upon salt stress, putrescine levels are elevated in *Oryza sativa* and *Nicotiana tabacum*, on the other hand, drought stress also induces putrescine in Arabidopsis and *Oryza sativa* (Roy & Wu, 2001; Kumria & Rajam, 2002; Alcázar *et al*., 2010; Capell *et al*., 2004). Putrescine biosynthetic mutants (*adc1* and *adc2*) in Arabidopsis with decreased levels of putrescine, resulted in altered responses against freezing (Cuevas *et al*., 2008). Role of putrescine is also known from other symbiotic interactions. In rice, a common metabolomic signature upon interaction with plant growth promoting rhizobacteria, is the increased accumulation of hydroxycinnamic acid amides (HCAA), identified as *N-p*-coumaroyl putrescine and *N*-feruloyl putrescine (Valette *et al*., 2019). An accumulation of coumaroyl putrescine, and N-feruloyl putrescine is also reported in early developmental stages of barley mycorrhization (Peipp *et al*., 1997). Putrescine when supplied exogenously induced the growth of *S. lycopersicum* and *P. indica* indicating that the polyamine is co-adapted by both host and microbes. In plants, it is known that low amounts of these exogenous polyamines can act as growth stimulants (Martin-Tanguy 2001). Homeostasis of polyamines is tightly regulated because higher levels of polyamines are toxic to cells and causes cell death (Kusano *et al*., 2008). For example, overexpression of polyamine biosynthesis gene *ADC2* reduced growth in Arabidopsis (Alcázar et al., 2005). Elevation of *in vitro* growth of *P. indica* upon exogenous putrescine treatment suggests its role in hyphal growth. Interestingly, higher concentration of putrescine than 10 μM leads to the saturation level of growth promotion, which indicates optimal concentration of putrescine for growth promotion. Putrescine and spermidine are involved in regulating hyphal growth of arbuscular mycorrhizal fungi, *Glomus mosseae* due to endogenous concentration of these compounds in spores being a growth limiting factor (Ghachtouli *et al*., 1996).

It is well known that PA interacts with hormones to regulate the growth and development of plants (Liu *et al*., 2013, Li *et al*., 2018). Auxin and cytokinin are major growth hormones involved in plant growth promotion response of *P. indica* in diverse plants (Xu *et al*., 2018). *P. indica* induces a rapid increase in auxin levels during early recognition phases which is crucial for reprogramming root development (Meents *et al*., 2019). Auxin level increased upon *P. indica* infestation in Arabidopsis roots (Vadassery *et al*.,2008) and *P. indica* also produces IAA in liquid culture (Sirrenberg *et al*., 2007). Several reports demonstrate that *P. indica* interferes with auxin production and signaling in the hosts and contribute to root growth (Sirrenberg *et al*., 2007; Vadassery *et al*., 2008; Lee *et al*., 2011; Dong *et al*., 2013; Ye *et al*., 2014; Kao *et al*., 2016; Hua *et al*., 2017). A significant increase in auxin (IAA) level in *S. lycopersicum* seedlings upon putrescine treatment confirmed putrescine-mediated auxin alteration. Interestingly, auxin responsive elements are located in the promoters of *SlADC1* (Liu *et al*., 2018) and could play a role in its regulation. *P. indica* colonization is known to up-regulate several GA biosynthesis genes. GA-deficient *ga1-6* mutant reduce *P. indica* colonization, whereas the quintuple-DELLA mutant can increase colonization (Schäfer et al., 2009; Jacobs et al., 2011). GA elevation in rice roots by *P. indica* can induce plant tolerance to root herbivory (Cosme et al., 2016). Furthermore, polyamines are observed to be involved in gibberellin induced development in grape berries and peas (Shiozaki et al., 1998; Smith et al., 1985). Therefore, increased level of GA_4_ and GA_7_ in putrescine treated *S. lycopersicum* seedlings might be crucial for growth promotion. How putrescine a primary metabolite involved in diverse plant process has been co-adapted by a symbiotic microbe for enhancing bi-directional growth is the focus of future study.

## Conclusion

*P. indica* colonization realigns the *S. lycopersicum* metabolism including polyamine biosynthetic pathway in root, where biosynthesis of a major polyamine, putrescine, is significantly induced through ADC-mediated pathway. Putrescine plays a vital role during interaction between *P. indica* and *S. lycopersicum* as well as Arabidopsis and is inevitable for their growth promotion by *P. indica*. Our work sheds the light on a mechanism where *P. indica* employs *ADC*-mediated putrescine biosynthesis in *S. lycopersicum* to promote its own growth into the host plant’s roots. Simultaneously, host plant allows induction of putrescine biosynthesis as it helps in plant’s growth promotion via Auxin and GA induction (Fig. 8).

**Figure 8.**
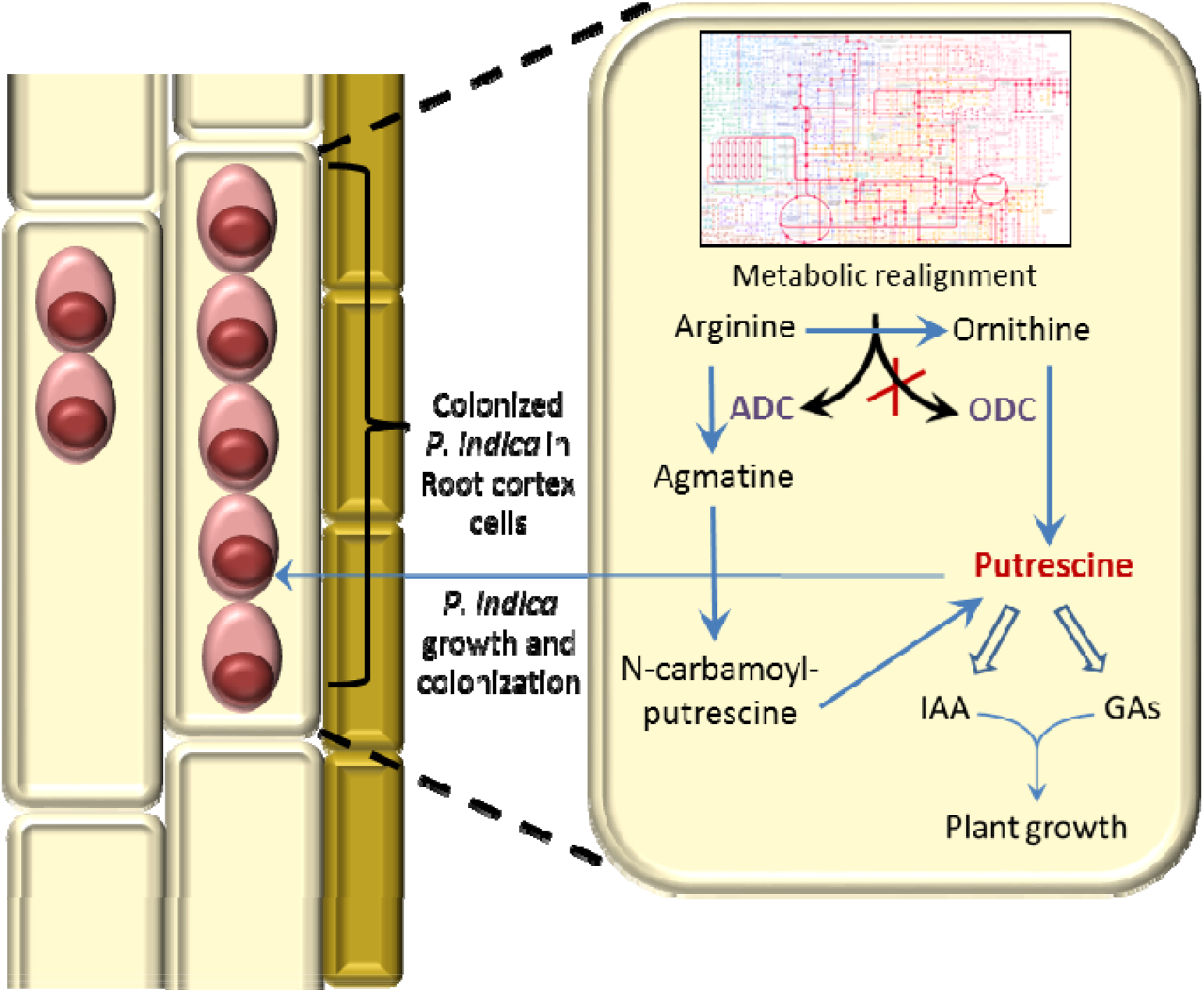
Schematic representation of *P. indica* induced putrescine biosynthesis in plants that promotes the growth of both plants and *P. indica*. *P. indica* realigns cellular metabolism and induces arginine decarboxylase (ADC) mediated putrescine biosynthesis. The other putrescine biosynthetic pathway mediated by ornithine decarboxylase (ODC) is not induced by *P. indica*. The increased biosynthesis of putrescine induces IAA and GAs which promotes growth in plants. Induced putrescine level also helps in *P. indica* growth.

## Materials and Methods

### Plant growth and *P. indica* co-cultivation

*P. indica* (Verma et al., 1998) culture was grown and maintained on Kaefer medium (Varma et al., 1999) at 28±2°C and 110 rpm in orbital shaker. Tomato (*Solanum lycopersicum*, cv. Pusa Ruby) seeds were pre-soaked in water overnight, kept on a moist tissue paper for 5 days in dark and after germination co-cultivated with 1% *P. indica* (w/w) in soilrite (horticulture grade expanded perlite, irish peat moss, and exfoliated vermiculite in equal ratio i.e., 1:1:1, w:w:w). The plants were grown at 26°C (day/night: 16/8 h), relative humidity 60% and light intensity 300 μmol/m^2^/ sec for different time points (10dpi, 20dpi, 30dpi and 40dpi). Control plants were cultivated without *P. indica* inoculation. Fungal colonization was detected using trypan blue staining of root segments (Fig. S1). Simultaneously, tomato roots colonized with GFP-tagged *P. indica* (Hilbert et al., 2012, Jogawat et al., 2020) were also harvested for visualization of colonization under confocal microscope (Leica TCS M5). GFP-tagged *P. indica* were received from Prof. Ralf Oelmüller (Friedrich Schiller University, Jena, Germany).

For experiments with Arabidopsis, seeds of wild type (ecotype Columbia) and *adc 1-2* (SALK_085350C), *adc 1-2* (CS9658), *adc 1-3* (CS9657) and *adc 2-3* (CS9659), *adc 2-4* (CS9660) mutant lines with T-DNA insertion in the exons (Alonso et al., 2003) from TAIR were used. Seeds were surface-sterilized, stratified and placed on half-strength MS plates supplemented with 1% sucrose and 0.8% agar and germinated for 7 days. The seedlings were grown at 22°C, 10 h light/14 h dark photoperiod and a light intensity of 150 μmol m^−2^ s^−1^ in growth chamber (Percival).These seedlings were transferred to 1× PNM (plant nutrient medium) medium (Hilbert et al., 2012, Johnson et al., 2013) for co-cultivation with *P. indica*. Samples were harvested at 14dpi stage

### Visualization and study of *P. indica* colonization

For observation of *P. indica* colonization in *S. lycopersicum*, the roots were harvested, washed and softened using 10% KOH, acidified in 1 M HCl and then stained with 0.02% Trypan blue for 1 h at 65°C. The samples were then de-stained in 50% lacto-phenol for 2 h (Dickson and Smith, 1998). The roots were observed under light microscope (Nikon 80i). For confocal microscopy, the tomato roots were colonized with GFP-tagged *P.indica* strain (Hilbert et al., 2012). At 40 dpi the roots were harvested and observed under a confocal microscope (Leica TCS M5) at an emission wavelength of 505-530 nm, 470 nm excitation and digital sectioning of 4-5 μm of root thickness (Jogawat et al., 2020).The relative amount of fungal DNA was quantified in 30 and 40 dpi roots using real time-qPCR utilizing *UBI3* and *P. indica Tef1* (Bütehorn et al., 2000). Relative changes in fungal DNA content were calculated using C_T_ of *PiTef1* which were normalized by C_T_ of *UBI3* using ΔΔCT equation and *P. indica* DNA content in control roots (0 dpi) was defined as 1.0 (Vadassery et al., 2008, Jogawat et al., 2020). The primer pairs used in the gene expression studies are given in Table S1.

### Untargeted metabolite profiling through GC-MS

Leaf (all the leaflets) and roots of *P. indica* colonized *S. lycopersicum* (40dpi) and *P. indica* mycelia were harvested and lyophilized. Three biological replicate, which was a pool of three individual samples were taken for each analysis. Lyophilized samples were extracted and derivatized for GC-MS analysis according to Kundu et al., 2018. The derivatized samples were analyzed through gas chromatography-mass spectrometry (GC-MS) by a Shimadzu GC-MS-QP2010TM coupled with an auto sampler-auto injector (AOC-20si). Chromatography was performed using an Rtx-5^®^ capillary column (Restek Corporation, US) and helium as carrier gas. Peak integration and mass spectra analysis were done through GC-MS solution software (Shimadzu^®^). Derivatized metabolites were identified through aligning and matching the mass spectra with NIST14s spectral library. Normalization of each peak area was done by internal standard’s (ribitol) peak area used in each sample. For some metabolites with multiple peaks, summation of the normalized peak area was considered after confirming the mass spectra as per the published protocol (Lisec et al., 2006). Venn diagram was generated in Venny 2.1 (https://bioinfogp.cnb.csic.es/tools/venny/). Logarithmic values of normalized peak area and metabolite fold change were used for all multivariate statistical analysis considering FDR. Volcano plots were prepared in Origin 6.0 (https://www.originlab.com/) by using Log_10_ values of the fold change (FC), where FC>1.5 was taken as cut-off value and *P*<0.05 was taken as cut-off for significance. Pearson’s correlation based clustered heat-map, PLS-DA, OPLS-DA, network analysis and pathway impact analysis were done by using MetaboAnalyst 4.0 (http://www.metaboanalyst.ca/). Metabolite network analysis was done by significantly altered metabolite fold changes and pathways were selected those have significance *P*<0.05. Correlation network was constructed by Cytoscape 3.2.0 aided with MetScape by uploading correlation values calculated in correlation algorithm of MetaboAnalyst 4.0.

### Putrescine quantification through LC-MS/MS

Extraction and derivatization through benzoylation of putrescine was done according to Lou et al., 2016 with slight modification. In brief, around 200-250 mg of fresh plant sample was ground in liquid nitrogen and extracted with 1 mL of 10% perchloric acid. The extract was centrifuged at 12000 g for 10 min at 4°C. Supernatant was collected and 500 μL of supernatant was derivatized with 500 μL of 2N sodium hydroxide and 20 μL of benzoyl chloride by incubating the mixture at 48°C for 20 min. 1 mL of sodium chloride was added to the sample and extracted with 1 mL of diethylether. Centrifugation was done at 5000 g for 10 min and the upper phase of the sample was collected and evaporated to dryness. The dried derivatized sample was again dissolved in 500 μL of 50% acetonitrile. This sample was diluted (1:1000, v/v) with 50% acetonitrile before analyzing it in LC-MS/MS. Derivatized putrescine was analyzed in Exion LC (SCIEX) coupled with triple-quadruple-trap MS/MS equipped with a Turbospray ion source (SCIEX 6500+).

Chromatography was performed on a Zorbax Eclipse XDB-C_18_ column (50 × 4.6 mm, 1.8 μm, Agilent Technologies) by using 1% formic acid (solvent A) and acetonitrile (solvent B) as mobile phase. A linear gradient (0-1 min, 5% B; 1-7 min, 5-95% B; 7-7.6 min, 95-5% B; 7.6-9 min, 5% B) was applied for derivatized putrescine separation. For detection, the mass spectrometer was operated in MRM mode to monitor analyte parent ion → product ion (297.0→105). Settings were as follows: ion spray voltage, 5500 eV; turbo gas temperature, 650°C; nebulizing gas, 70 p.s.i.; curtain gas, 45 p.s.i.; heating gas, 60 p.s.i.; DP, 60; EP,10; CE, 20; CXP,10. Quantification was done by using external calibration curve prepared with authentic putrescine standard (Sigma^®^).

### Growth phytohormone estimation

250 mg of *S. lycopersicum* seedlings were ground in liquid nitrogen and extracted with 1 mL of cold extraction buffer (MeOH: H_2_O:HCOOH, 15:4:0.1) containing 25 ng of *trans*-[^2^H_5_] zeatin, *trans*-[^2^H_5_] zeatin riboside, [^2^H_6_] N^6^-isopentenyladenine, [^2^H_5_]-indole-3-acetic acid (^2^H_5_-IAA) and [^2^H_2_]-GA_1_ as internal standards. Homogenized sample was centrifuged at 10,000 g for 10 min at 4°C. Supernatant was loaded onto a Strata-X (Phenomenex ^®^) C_18_ solid phase extraction (SPE) column pre-conditioned with 1 mL of methanol and 1 mL of 0.1% formic acid in water. After loading, the SPE column was washed twice with 0.1% formic acid and 5% methanol. Finally, the elution was done with 1 mL 0.1% formic acid in acetonitrile and dried in speed-vac. Dried sample was re-dissolved in 100 μL 5% methanol and analyzed by liquid chromatography coupled with a SCIEX 6500^+^ triple-quadruple-trap MS/MS. LC-MS/MS was performed according to (Schäfer *et al*., 2013) with slight modifications. In brief, separation of phytohormone was done with a Zorbax Eclipse XDB C18 column (50 × 4.6 mm, 1.8 μm, Agilent Technologies) was used. The mobile phase comprised solvent A (water, 0.05% formic acid) and solvent B (acetonitrile) with the following elution profile: 0-0.5 min, 95% A; 0.5-5 min, 5-31.5% B in A; 5.01-6.5 min 100% B and 6.51-9 min 95% A, with a flow rate of 1.1 mL min^−1^. The column temperature was maintained at 25°C. For detection of IAA and cytokinins, the mass spectrometer was operated in positive ionization mode (MRM modus) to monitor analyte parent ion → product ion (176.0→130.0 for IAA; 220.2→136.3 for *trans*-zeatin; 352.2→220.3 for *trans*-zeatine riboside, 354.2→222.1 for dihydrozeatin riboside, 514.1→382.1 for *trans*-zeatin riboside-O-glucoside, 382.1→220.19 for *trans*-zeatin-7-glucoside, 222→136 for dihydrozeatin, 516.2→222 for dihydrozeatin riboside-*O*-glucoside, 384.2→222 dihydrozeatin-*O*-glucoside, 225.2→136.3 for *trans*-[^2^H_5_]zeatin; *trans*-[^2^H_5_]zeatin riboside, 204.1→136 for isopentenyladenine, 210.1→136 for [^2^H_6_] N6-isopentenyladenine, 181.0→134.0 for [^2^H_5_]-IAA). Settings were as follows: ion spray voltage, 5500 eV; turbo gas temperature, 650°C; nebulizing gas, 70 p.s.i.; curtain gas, 45 p.s.i.; heating gas, 60 p.s.i. Analyst 1.5 software (Applied Biosystems) was used for data acquisition and processing. For detection of GAs, MS analysis triple Quad 6500+ is operated in negative ionization mode with Ion Spray Voltage of −4500 eV, CUR gas 45 psi, CAD-medium, Temperature 650, GS1 and GS2 60 psi. Multiple-reaction monitoring (MRM) is used to monitor analyte parent ion□→□product ion (331.1→213.1 for GA_4_, 345.1→143.1 for GA_3_, 347.1→273.1 for GA_1_, 329.1→223.1 for GA_7_, 363.1→275.1 for GA_8_ and 349.0→276.0 for [^2^H_2_]GA_1_) with detection window of 60 seconds.

### Preparation of virus induced gene silencing (VIGS) construct for *SlADC1* gene silencing

To knock-down the expression of *ADC* in *Solanum lycopersicum*, the TRV-VIGS technique was used as previously reported (Lee *et al*., 2017). The sequence of *SlADC1* gene (2124 bp) was obtained from the genome version: *Solanum lycopersicum* ITAG V3.2. For the *SlADC1-VIGS* silencing construct, a short sequence corresponding to 101 bp to 476 bp in the CDS sense strand was selected by SGN-VIGS TOOL (vigs.solgenomics.net/). The 375 bp short sequence was amplified and cloned in the TRV2 vector. Details of the primer pair is provided in Table S1. The silencing ability of the TRV2 vector was confirmed by the knock down of *Phytoene Desaturase* gene (*PDS*) gene responsible for chlorophyll biosynthesis in *S. lycopersicum*. The positive plasmids for TRV2*: SlADC* was confirmed by sequencing and transformed in the *Agrobacterium tumefaciens* strain *GV3101*. VIGS in *S. lycopersicum* was carried out as described (Senthil-Kumar and Mysore, 2014) with slight modifications by the method of needleless syringe inoculation (Ryu *et al*., 2004). Briefly, pTRV1, pTRV2 and pTRV2-*Sl*ADC1 constructs were grown till OD_600_=1.0 separately and mixed to 1:1 ratio before infiltration to the abaxial leaf surface of 14 days old plants with a 1 ml needle-less syringe for two independent sets of plants (TRV:00, and TRV-*SlADC1*) and were maintained in a plant growth room at 26°C (day/night: 16/8 h). At 7 and 14 days post infiltration (dpi), the infiltrated leaves were harvested for RNA isolation and cDNA synthesis. The relative transcript abundance of the *ADC1* gene was analyzed. For *P. indica-tomato* experiments, 14 days old plants post germination was co-cultivated with *P. indica* spores by applying 1 ml of spore suspension at a concentration of 5 x 10^5^ spores/ml at the crown region of the plant in soilrite simultaneously during agro-inoculation and were allowed to grow for 40 days.

### RNA extraction and gene expressions analysis

Harvested plant samples were homogenized using liquid N_2_ and total RNA was extracted using TRIzol Reagent (Invitrogen). Extracted RNA was treated with DNase (TURBO DNase, Ambion) to remove DNA contamination, and its quantification was done using Nano Drop. cDNA was prepared by using High capacity cDNA reverse transcriptase kit (Applied Biosystems^®^). Gene sequences were availed from Sol Genomics Network (https://solgenomics.net/), and gene-specific primers were designed using NCBI primer designing tool (https://www.ncbi.nlm.nih.gov/tools/primer-blast/). PowerUp™ SYBR green Master Mix (Applied Biosystems^®^) was used for the generation of amplicon. qRT-PCR was performed on a Bio-Rad CFX connect Real Time PCR. Relative expressions of the genes in the treated samples were calculated as fold change relative to untreated samples. Normalization of the gene expression was done with ubiquitin (*UBI3*) expression. The primer pairs used in the gene expression studies are given in Table S1.

### Putrescine and DFMA treatment on *P. indica* and plants

For examining the effect of putrescine on *P. indica* growth in axenic culture, five different concentrations (5, 10, 20, 50 and 100 μM) of putrescine (Sigma®) was directly supplemented to fungal defined minimal medium (MN medium) (Jogawat *et al*., 2016) and the fungus was allowed to grow at 28±2°C in the incubator. After 2 weeks, fungal radial growth and fresh weight was measured. Five-day-old seedlings of *S. lycopersicum* and *A. thaliana* grown in sterile conditions were transferred to solid half MS plates with 10 μM of putrescine and allowed to grow for 21 days. In a separate experiments five-day-old *S. lycopersicum* seedlings were transferred to half MS with the arginine decarboxylase inhibitor, DL-α-(Difluoromethyl) arginine (DFMA) (Santa Cruz Biotechnology ®) of 200 nM concentration and allowed to grow for 21 days.

### Statistical analysis

Significance analysis (T-test and One-way ANOVA) was done by using ‘Sigma Plot’, version 13 (www.sigmaplot.com). Plots of the figures were generated by using Origin 6.0 (www.originlab.com). Pearson correlation co-efficient was calculated by using ‘Social Science Statistics’ (https://www.socscistatistics.com/tests/pearson/) online tool and MetaboAnalyst 4.0.

## Supporting information

Chlamydospores were visible in the root cortex at 10 dpi with P. indica (Fig. S1)

The primer pairs used in the gene expression studies are given in Table S1.

55 were identified and annotated in leaf, 70 in root and 102 in P. indica mycelia (Fig. 2A, Table S2)

22 metabolites are shared by root, leaf and mycelia (Fig. 2B, Table S3).

GC-MS data-set for root and leaf specific metabolites (Table S4)

Pathway analysis with significantly altered (P<0.05) annotated root metabolite data set showed a plausible involvement of 46 metabolic processes (Fig.

Pathway analysis with significantly altered (P<0.05) annotated root metabolite data set showed a plausible involvement of 46 metabolic processes

## Acknowledgements

We acknowledge Department of Biotechnology (DBT), India for NIPGR core grant, and Max Planck-India partner group program of the Max Planck Society (Germany) for funding this work. We acknowledge JNU advanced instrumentation facility (AIRF) for GC-MS, NIPGR Metabolome facility funded by DBT (BT/ INF/22/SP28268/2018) for phytohormone quantification and LC-MS/MS analysis, Khushboo Adhlaka (Sciex, Gurgaon) for help in LC-MS/MS methods. We acknowledge NIPGR central instrumentation and phytotron facility and DBT-eLibrary Consortium (DeLCON) for providing access to e-resources. We also acknowledge Dr. Senthil Kumar Muthappa (NIPGR), New Delhi for providing VIGS vectors.

## Brief Legends for Supplemental Figures

**Supplemental Fig. S1** - Trypan blue staining shows *P. indica* colonization in *S. lycopersicum* root after 10 days of co-cultivation.

**Supplemental Fig. S2** - Different phenotypic changes and growth induction of *S. lycopersicum* upon *P. indica* colonization.

**Supplemental Fig. S3 –** Pearson’s correlation of metabolite alteration in leaf and root.

**Supplemental Fig. S4** - Alteration of leaf metabolites in *S. lycopersicum* upon *P. indica* treatment.

**Supplemental Fig. S5** – *S. Lycopersicum* root metabolites alteration upon *P. indica* colonization (40 dpi).

**Supplemental Fig. S6** - Scatter plot of pathway impact in root shows specific metabolism in root affected by *P. indica* colonization.

**Supplemental Fig. S7** - Effect of 10 μM putrescine on growth of *S. lycopersicum*.

**Supplemental Fig. S8** - Effect of Putrescine on *P. indica*

**Supplemental Fig. S9** - Schematic representation of the VIGS protocol in *Solanum lycopersicum*.

**Supplemental Fig. S10** - VIGS confirmation in *S. lycopersicum*.

**Supplemental Fig. S11**- *S. lycopersicum* and *P. indica* growth assay growth assay upon DL-α- (Difluoromethyl) arginine (DFMA) treatment.

**Supplemental Fig. S12**- Importance of putrescine on shoot and root growth of *A. thaliana* during *P. indica* infestation.

## Tables

**Supplemental Table S1.** Primer list.

**Supplemental Table S2.** Annotated metabolites in tomato and *P. indica* mycelia with derivatization level and obtained molecular mass.

**Supplemental Table S3.** Identity of metabolites distributed in Venn’s Diagram.

**Supplemental Table S4.** Fold change of metabolites found both in control and treated leaf and root samples.

**Supplemental Table S5.** Pathway impact analysis table for *S. lycopersicum* root treated with *P. indica*.

**Supplemental Table S6.** Metabolite-metabolite interaction analysis output.

