## Supplementary material for "*Piriformospora indica* employs host’s putrescine for growth promotion in plants": Chlamydospores were visible in the root cortex at 10 dpi with P. indica (Fig. S1)

### Slide 1
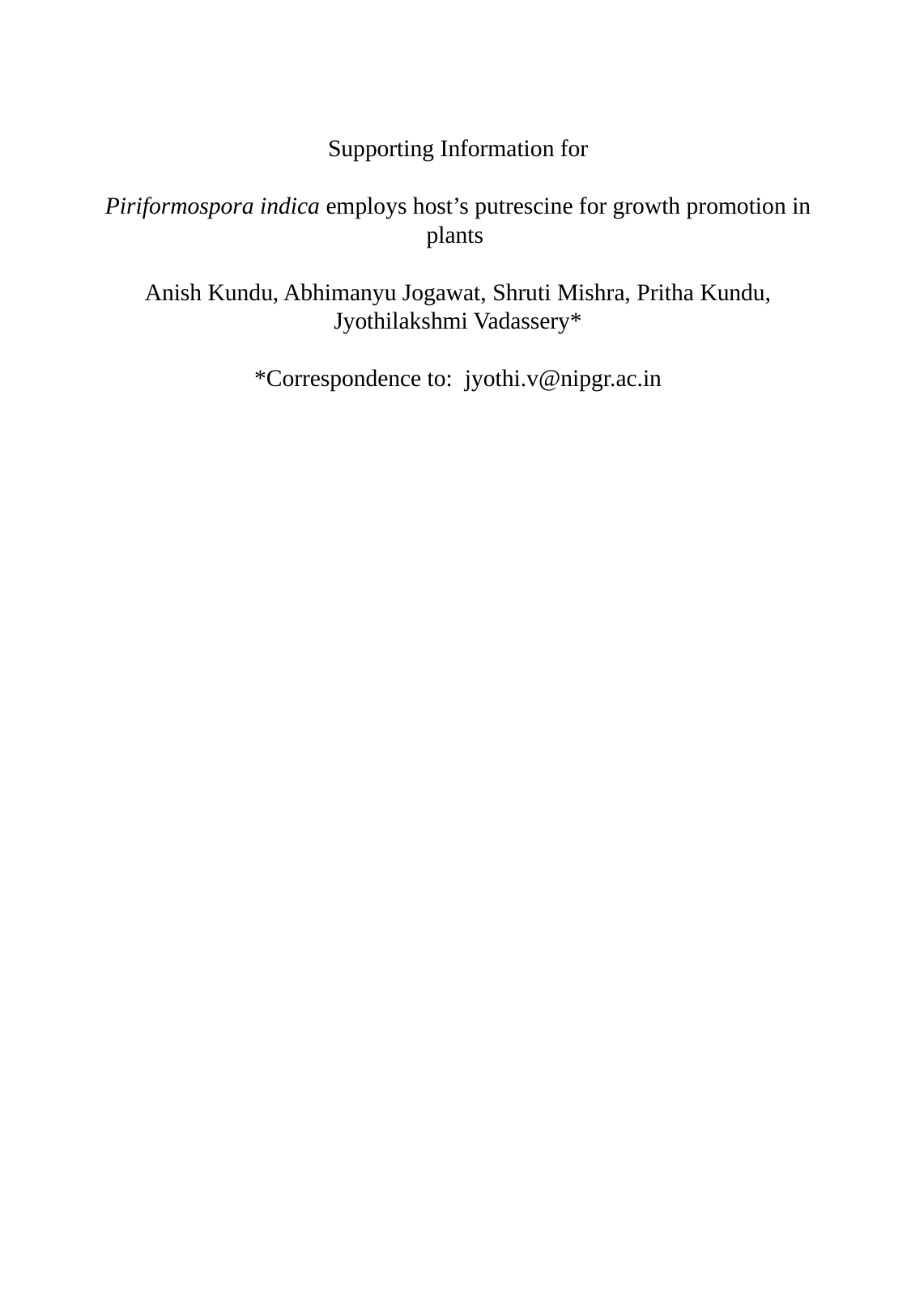

#### Supporting Information for Piriformospora indica employs host’s putrescine for growth promotion in plants Anish Kundu, Abhimanyu Jogawat, Shruti Mishra, Pritha Kundu, Jyothilakshmi Vadassery* *Correspondence to:

### Slide 2
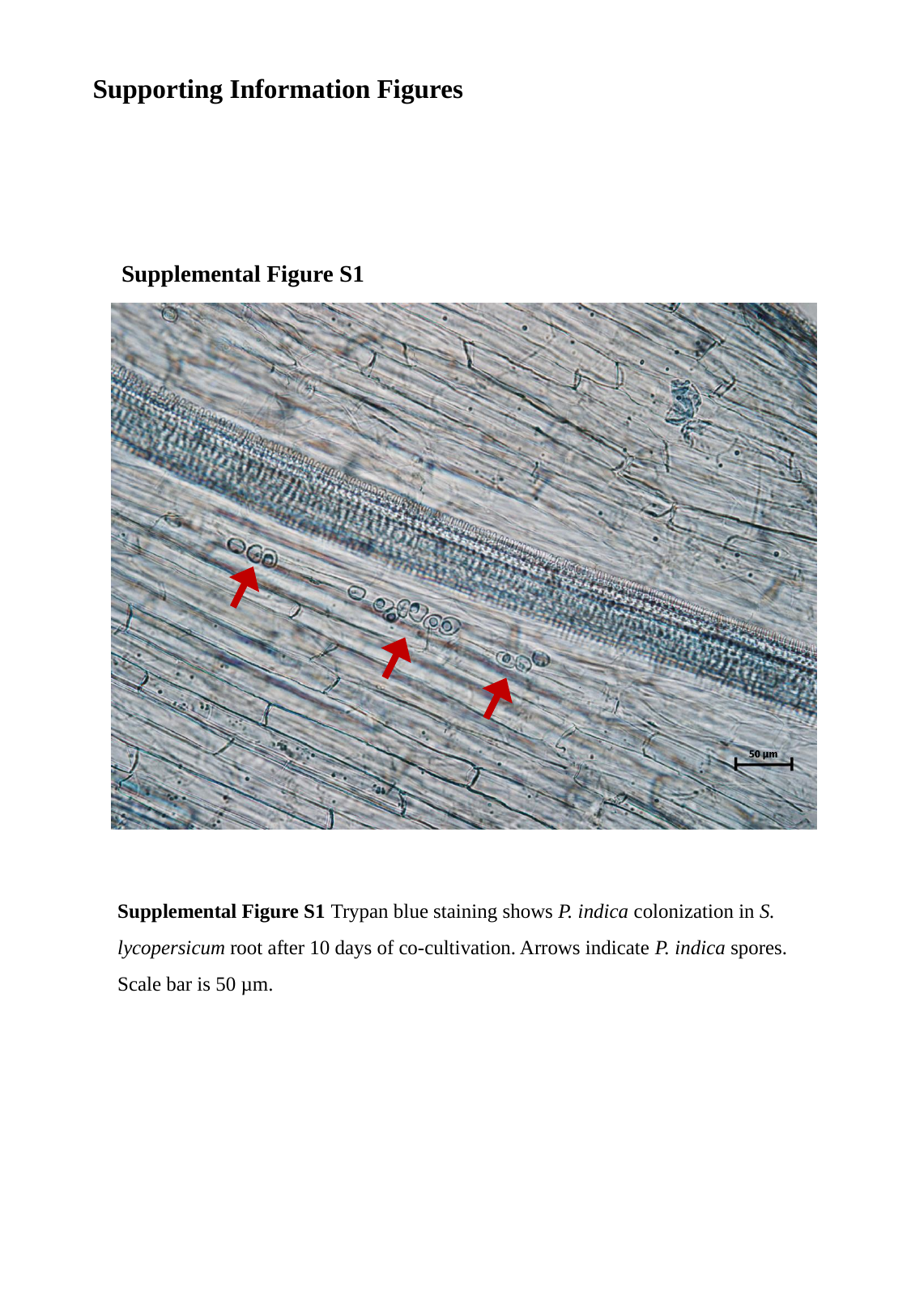

Supporting Information Figures
Supplemental Figure S1
Supplemental Figure S1 Trypan blue staining shows P. indica colonization in S. lycopersicum root after 10 days of co-cultivation. Arrows indicate P. indica spores. Scale bar is 50 µm.

### Slide 3
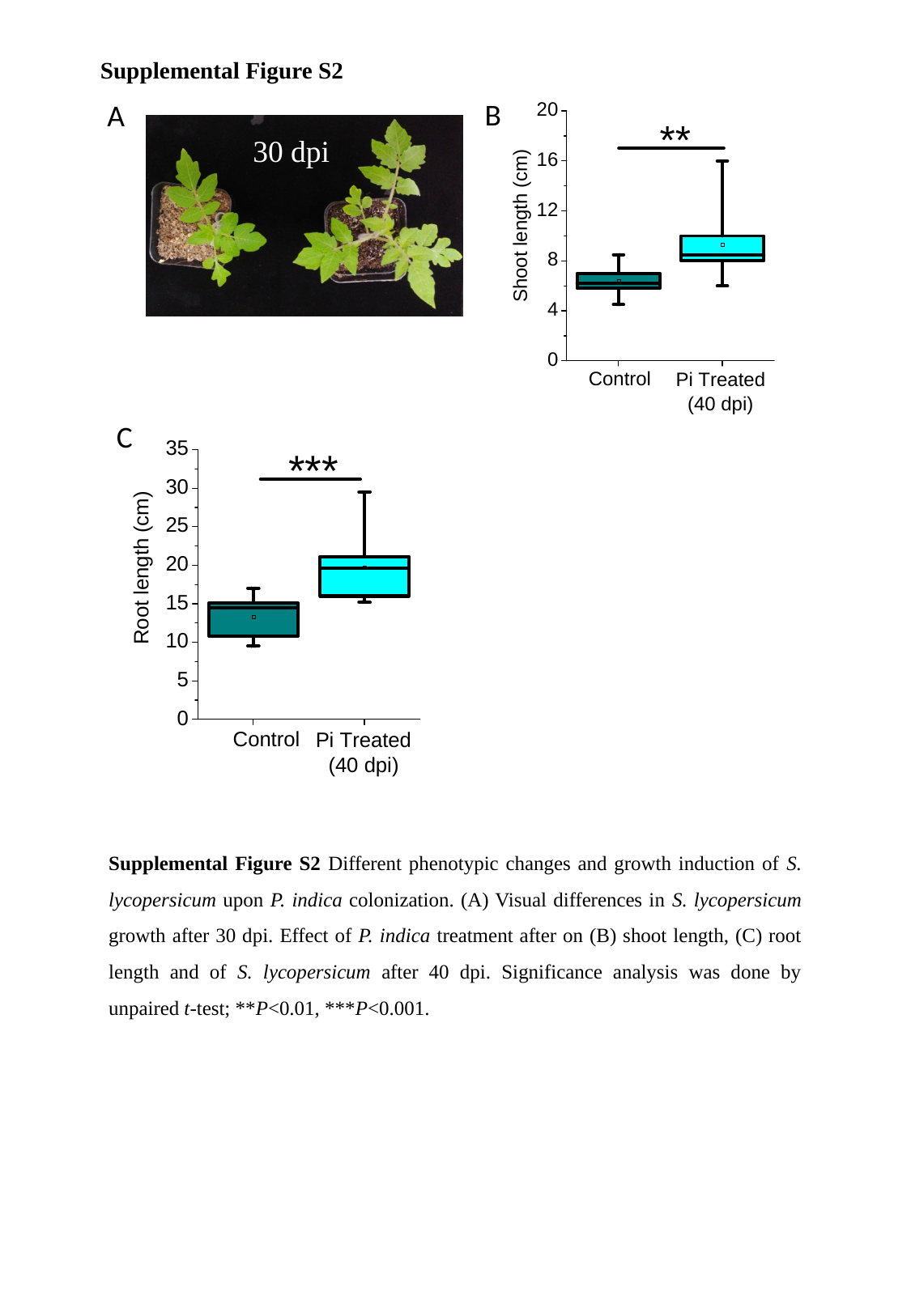

Supplemental Figure S2
A
30 dpi
Supplemental Figure S2 Different phenotypic changes and growth induction of S. lycopersicum upon P. indica colonization. (A) Visual differences in S. lycopersicum growth after 30 dpi. Effect of P. indica treatment after on (B) shoot length, (C) root length and of S. lycopersicum after 40 dpi. Significance analysis was done by unpaired t-test; **P<0.01, ***P<0.001.
B
C

### Slide 4
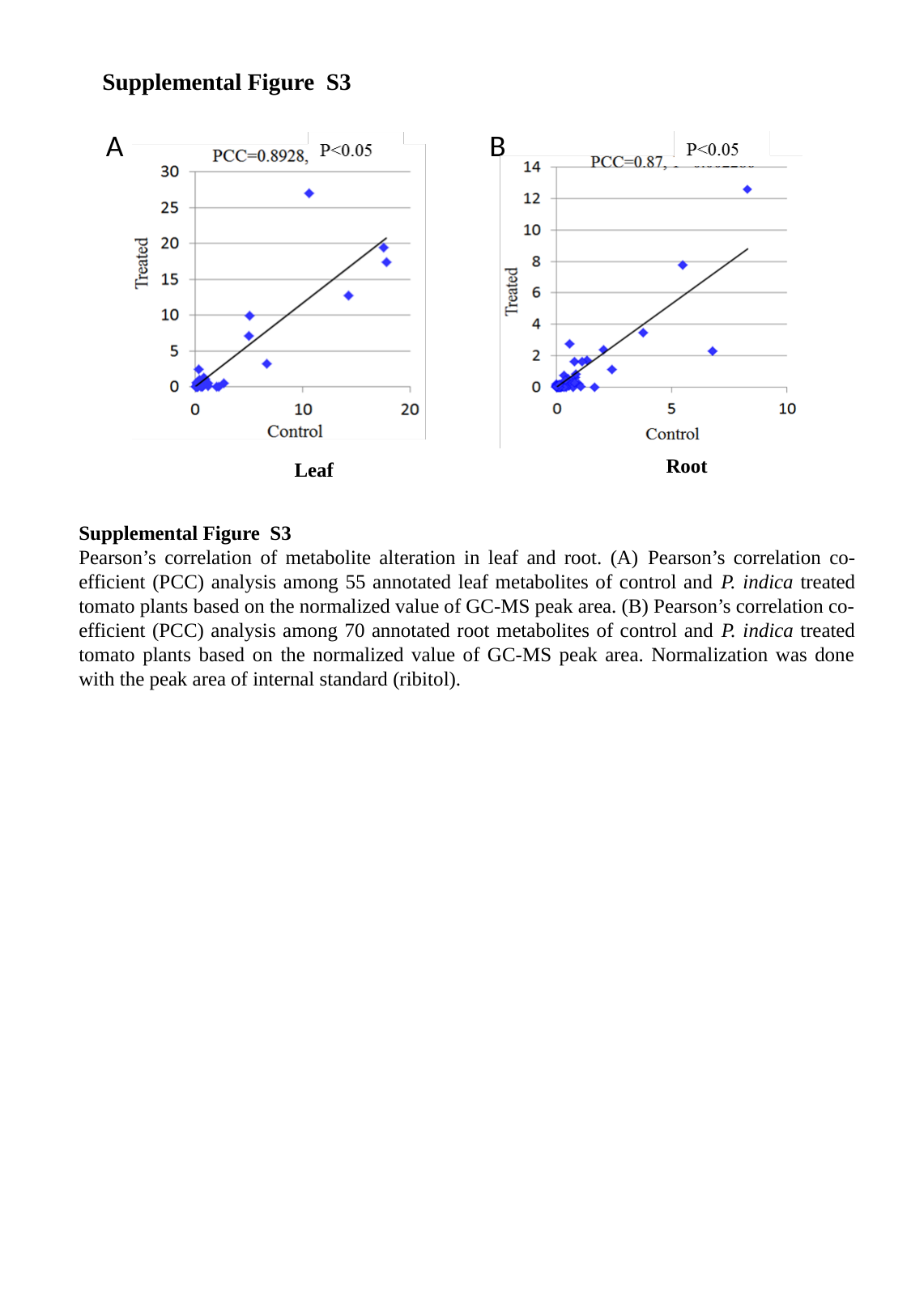

Supplemental Figure S3
A
B
Root
Leaf
Supplemental Figure S3
Pearson’s correlation of metabolite alteration in leaf and root. (A) Pearson’s correlation co-efficient (PCC) analysis among 55 annotated leaf metabolites of control and P. indica treated tomato plants based on the normalized value of GC-MS peak area. (B) Pearson’s correlation co-efficient (PCC) analysis among 70 annotated root metabolites of control and P. indica treated tomato plants based on the normalized value of GC-MS peak area. Normalization was done with the peak area of internal standard (ribitol).

### Slide 5
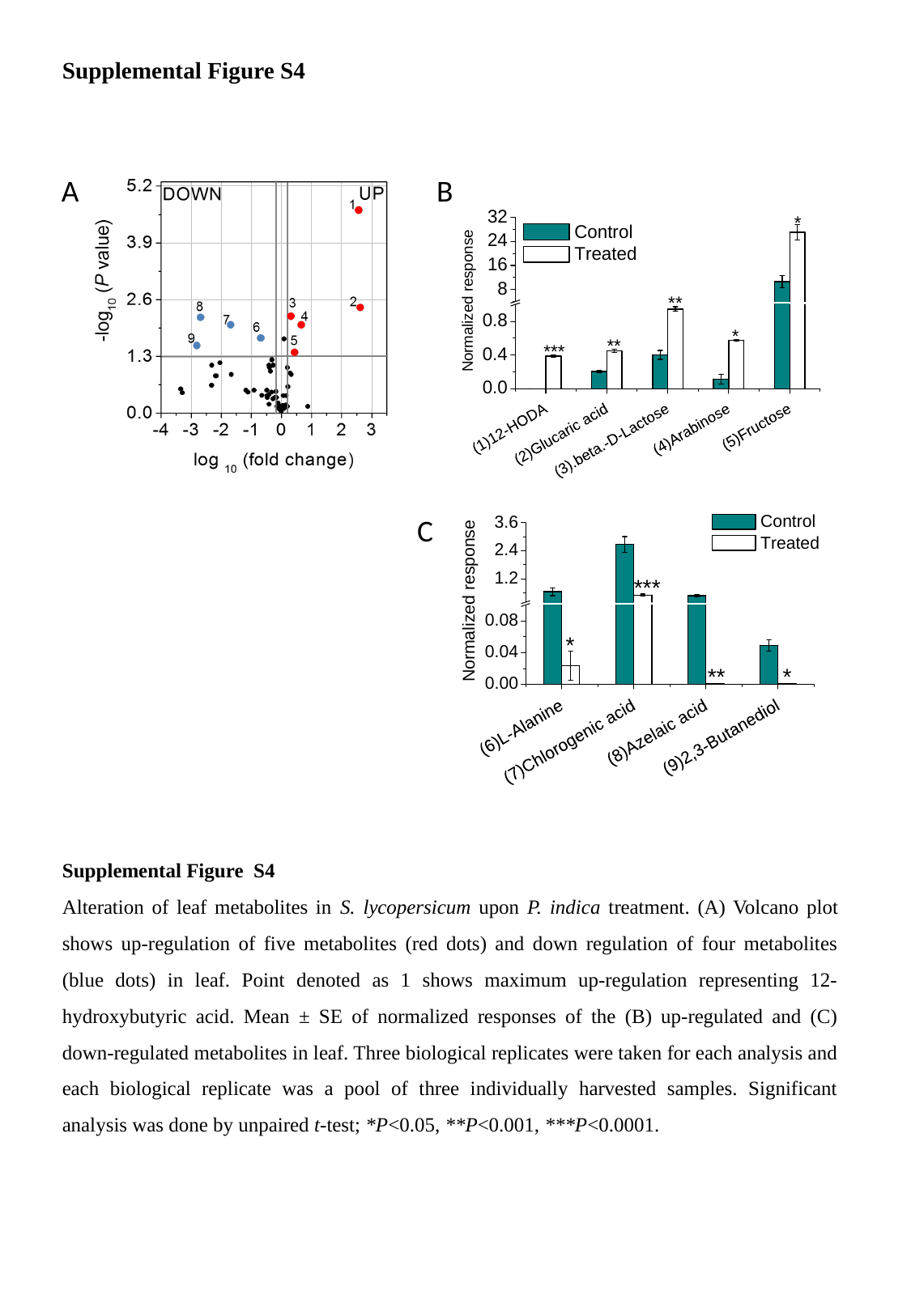

Supplemental Figure S4
A
B
C
Supplemental Figure S4
Alteration of leaf metabolites in S. lycopersicum upon P. indica treatment. (A) Volcano plot shows up-regulation of five metabolites (red dots) and down regulation of four metabolites (blue dots) in leaf. Point denoted as 1 shows maximum up-regulation representing 12-hydroxybutyric acid. Mean ± SE of normalized responses of the (B) up-regulated and (C) down-regulated metabolites in leaf. Three biological replicates were taken for each analysis and each biological replicate was a pool of three individually harvested samples. Significant analysis was done by unpaired t-test; *P<0.05, **P<0.001, ***P<0.0001.

### Slide 6
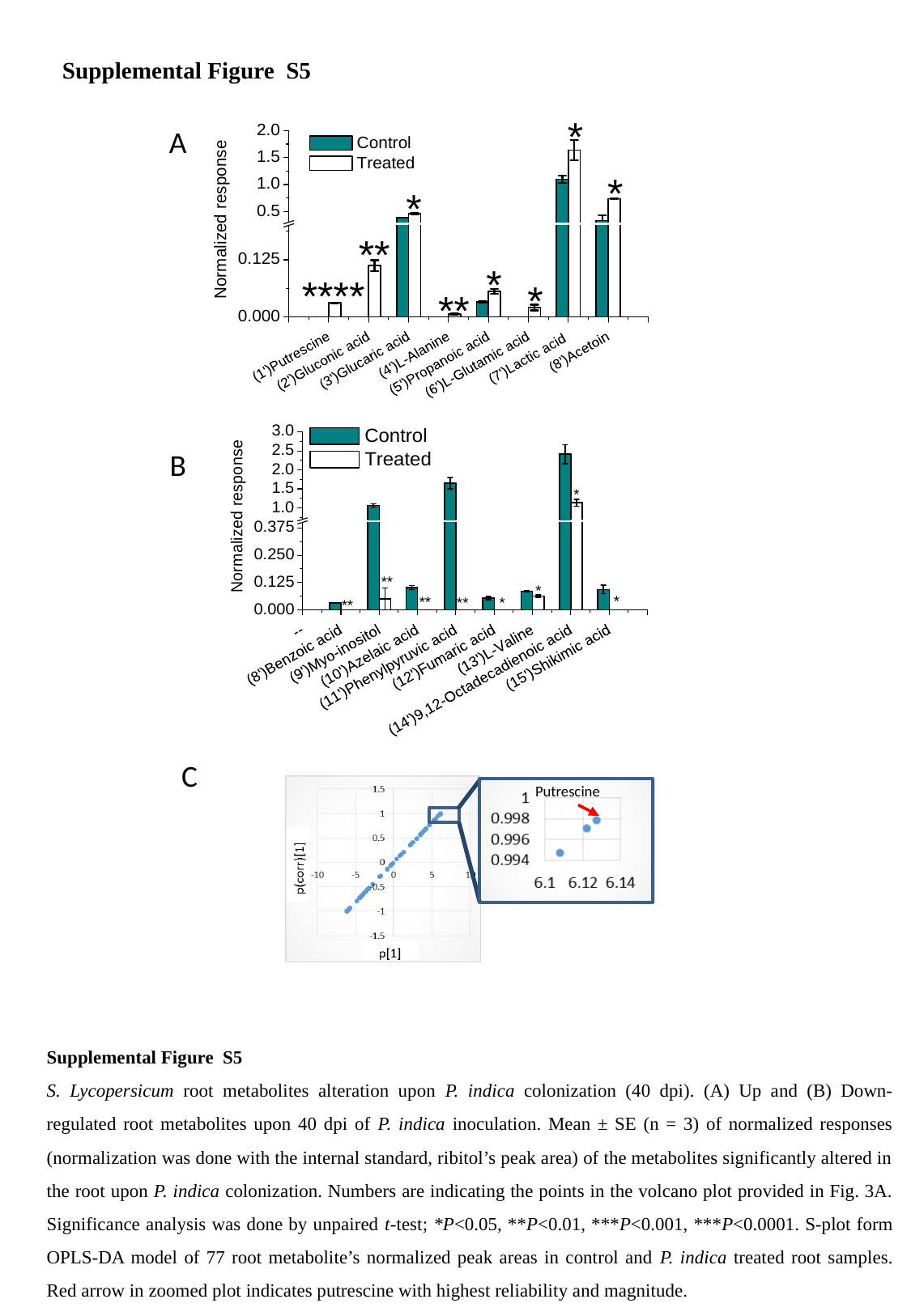

Supplemental Figure S5
A
B
C
Putrescine
Supplemental Figure S5
S. Lycopersicum root metabolites alteration upon P. indica colonization (40 dpi). (A) Up and (B) Down-regulated root metabolites upon 40 dpi of P. indica inoculation. Mean ± SE (n = 3) of normalized responses (normalization was done with the internal standard, ribitol’s peak area) of the metabolites significantly altered in the root upon P. indica colonization. Numbers are indicating the points in the volcano plot provided in Fig. 3A. Significance analysis was done by unpaired t-test; *P<0.05, **P<0.01, ***P<0.001, ***P<0.0001. S-plot form OPLS-DA model of 77 root metabolite’s normalized peak areas in control and P. indica treated root samples. Red arrow in zoomed plot indicates putrescine with highest reliability and magnitude.

### Slide 7
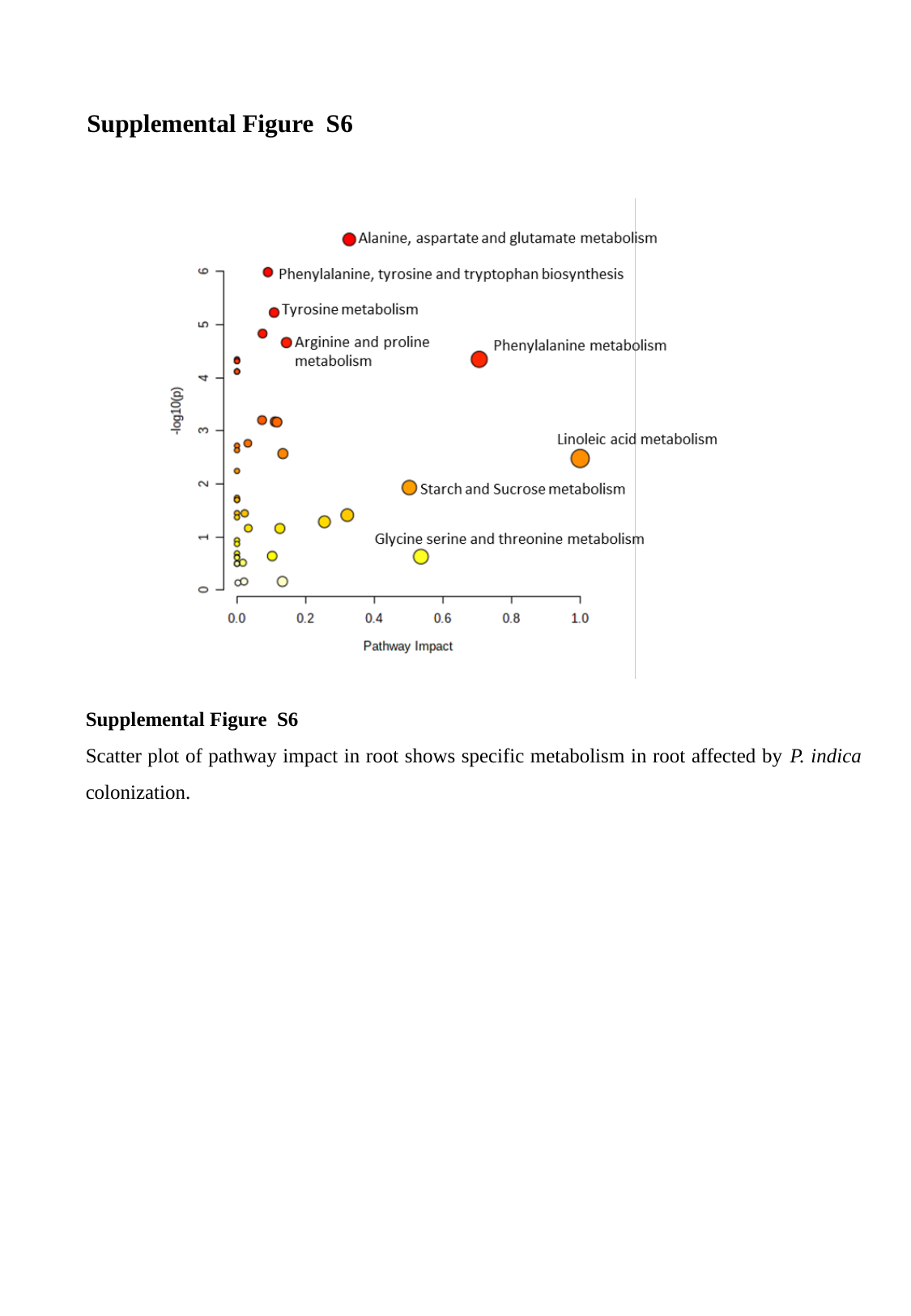

Supplemental Figure S6
Supplemental Figure S6
Scatter plot of pathway impact in root shows specific metabolism in root affected by P. indica colonization.

### Slide 8
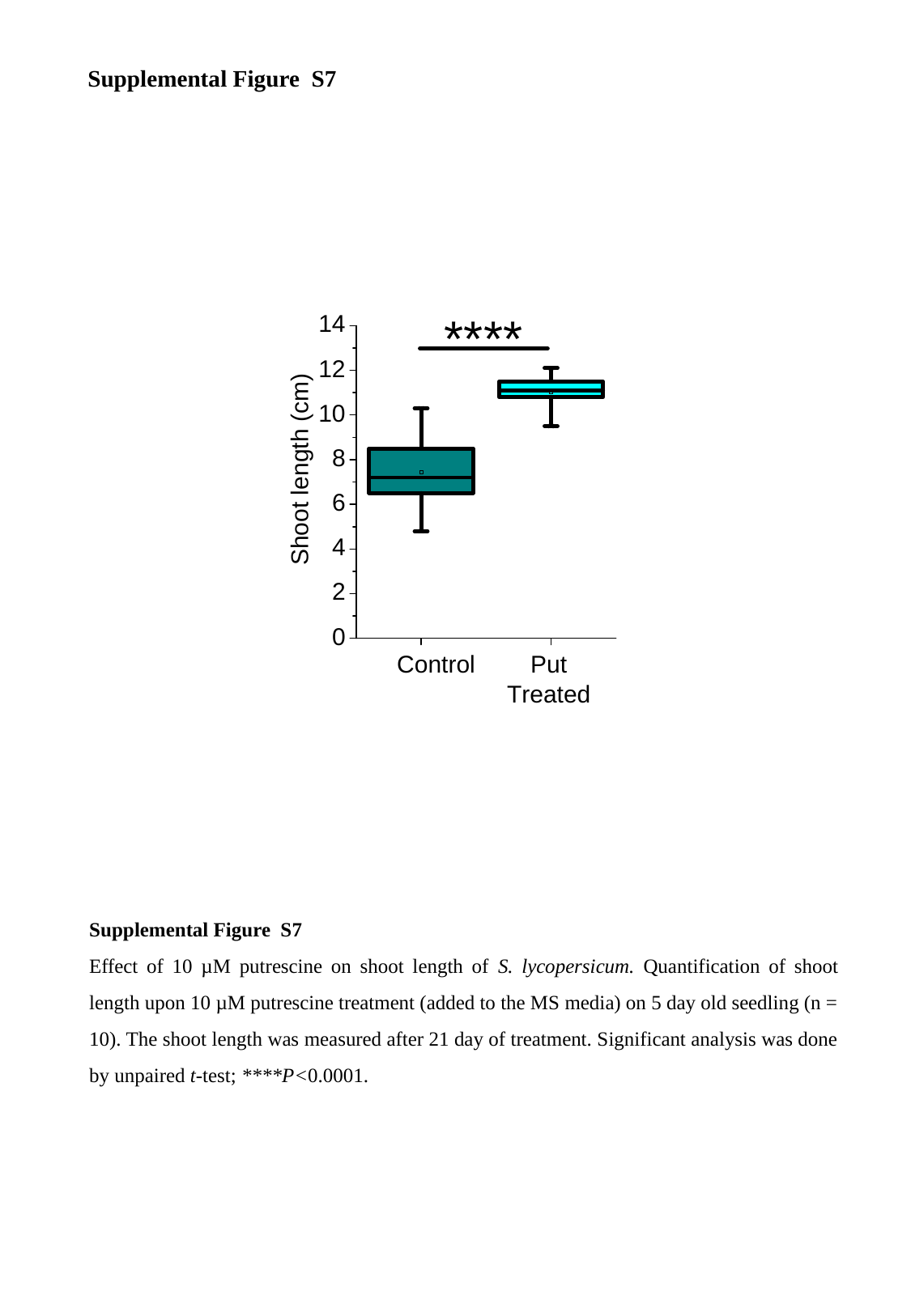

Supplemental Figure S7
Supplemental Figure S7
Effect of 10 µM putrescine on shoot length of S. lycopersicum. Quantification of shoot length upon 10 µM putrescine treatment (added to the MS media) on 5 day old seedling (n = 10). The shoot length was measured after 21 day of treatment. Significant analysis was done by unpaired t-test; ****P<0.0001.

### Slide 9
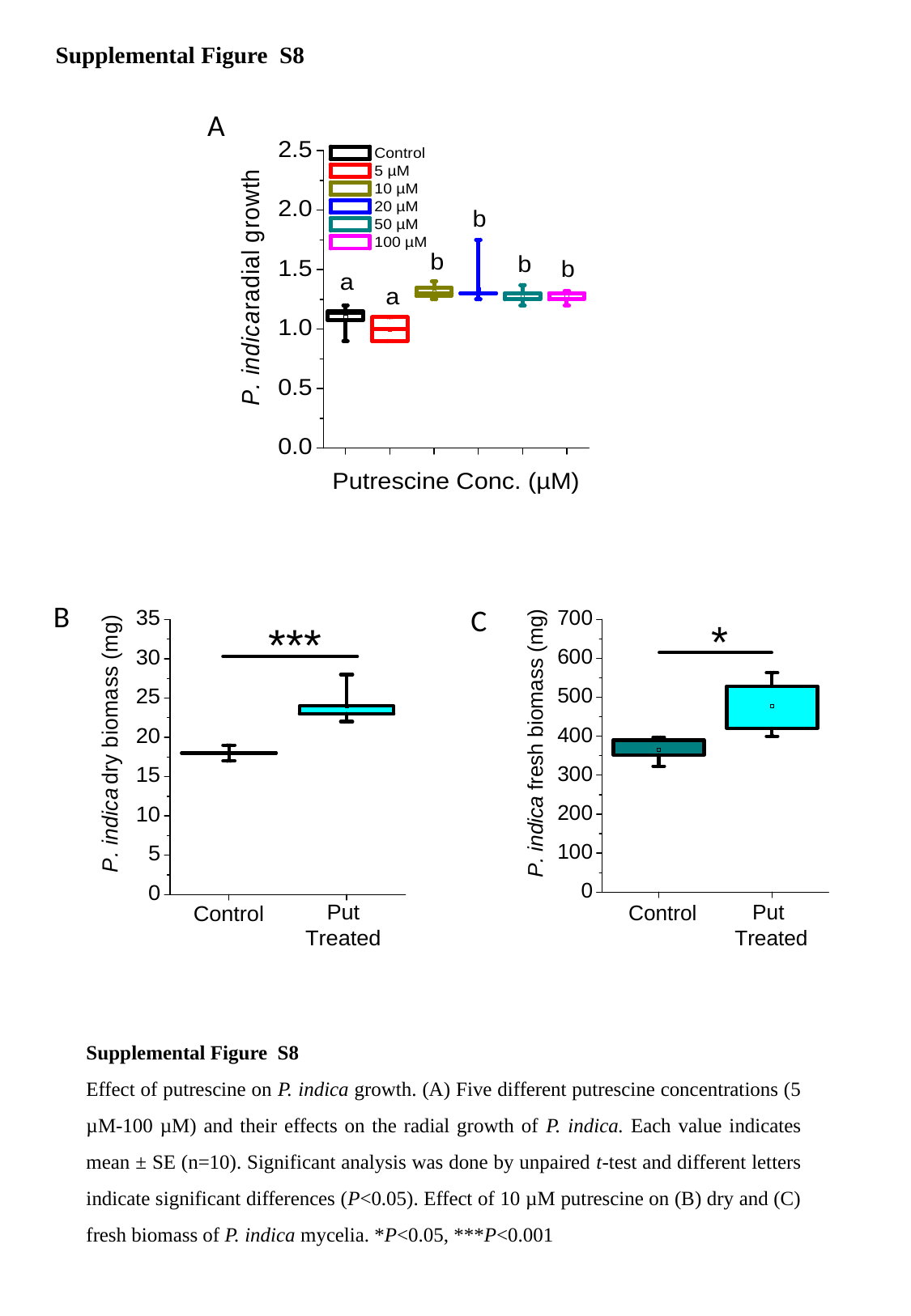

Supplemental Figure S8
A
B
C
Supplemental Figure S8
Effect of putrescine on P. indica growth. (A) Five different putrescine concentrations (5 µM-100 µM) and their effects on the radial growth of P. indica. Each value indicates mean ± SE (n=10). Significant analysis was done by unpaired t-test and different letters indicate significant differences (P<0.05). Effect of 10 µM putrescine on (B) dry and (C) fresh biomass of P. indica mycelia. *P<0.05, ***P<0.001

### Slide 10
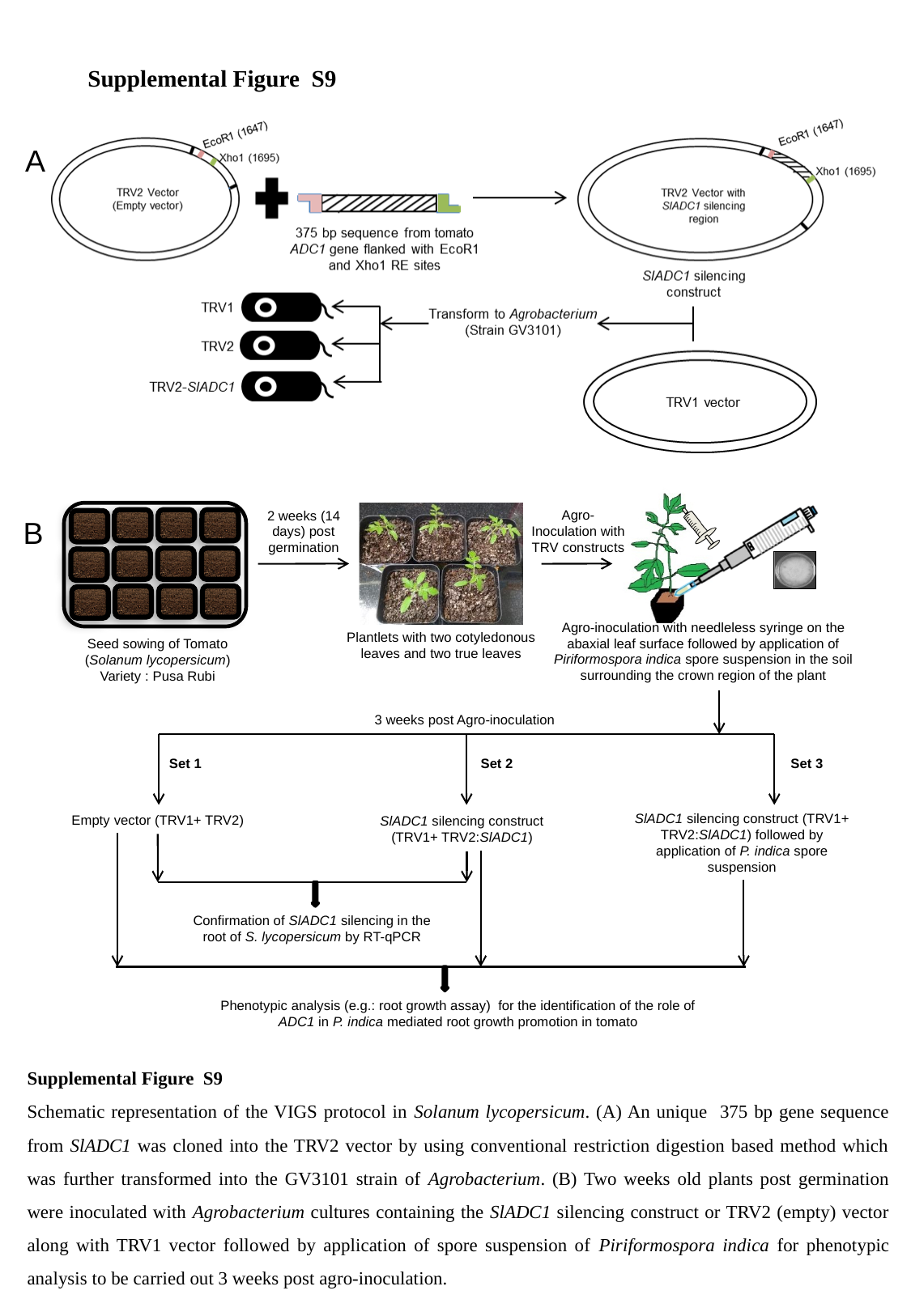

Supplemental Figure S9
A
Agro-Inoculation with TRV constructs
2 weeks (14 days) post germination
Agro-inoculation with needleless syringe on the abaxial leaf surface followed by application of Piriformospora indica spore suspension in the soil surrounding the crown region of the plant
Plantlets with two cotyledonous leaves and two true leaves
Seed sowing of Tomato (Solanum lycopersicum) Variety : Pusa Rubi
3 weeks post Agro-inoculation
Set 1
Set 2
Set 3
SlADC1 silencing construct (TRV1+ TRV2:SlADC1) followed by application of P. indica spore suspension
Empty vector (TRV1+ TRV2)
SlADC1 silencing construct (TRV1+ TRV2:SlADC1)
Confirmation of SlADC1 silencing in the root of S. lycopersicum by RT-qPCR
Phenotypic analysis (e.g.: root growth assay) for the identification of the role of ADC1 in P. indica mediated root growth promotion in tomato
B
Supplemental Figure S9
Schematic representation of the VIGS protocol in Solanum lycopersicum. (A) An unique 375 bp gene sequence from SlADC1 was cloned into the TRV2 vector by using conventional restriction digestion based method which was further transformed into the GV3101 strain of Agrobacterium. (B) Two weeks old plants post germination were inoculated with Agrobacterium cultures containing the SlADC1 silencing construct or TRV2 (empty) vector along with TRV1 vector followed by application of spore suspension of Piriformospora indica for phenotypic analysis to be carried out 3 weeks post agro-inoculation.

### Slide 11
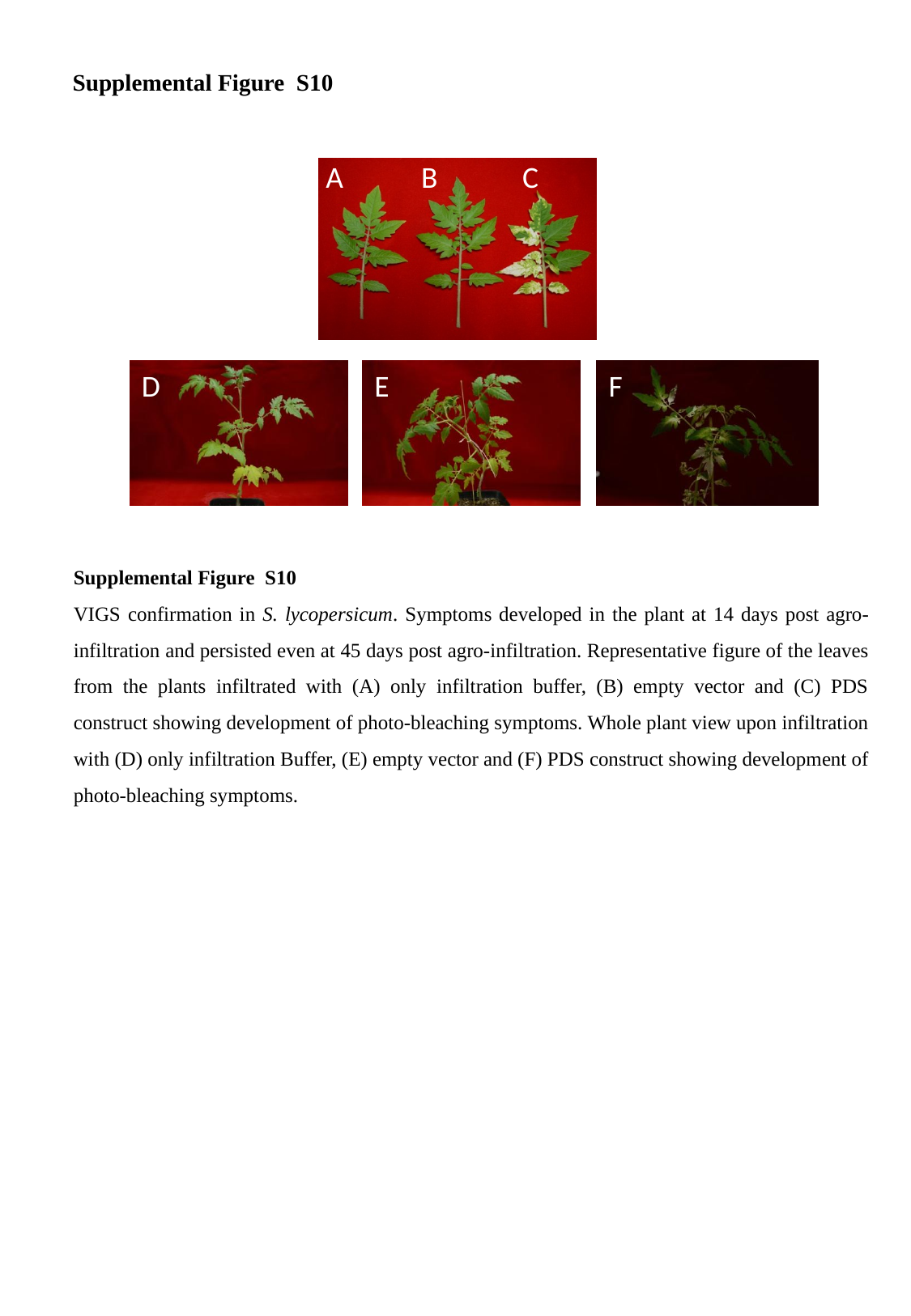

Supplemental Figure S10
A
B
C
E
D
F
Supplemental Figure S10
VIGS confirmation in S. lycopersicum. Symptoms developed in the plant at 14 days post agro-infiltration and persisted even at 45 days post agro-infiltration. Representative figure of the leaves from the plants infiltrated with (A) only infiltration buffer, (B) empty vector and (C) PDS construct showing development of photo-bleaching symptoms. Whole plant view upon infiltration with (D) only infiltration Buffer, (E) empty vector and (F) PDS construct showing development of photo-bleaching symptoms.

### Slide 12
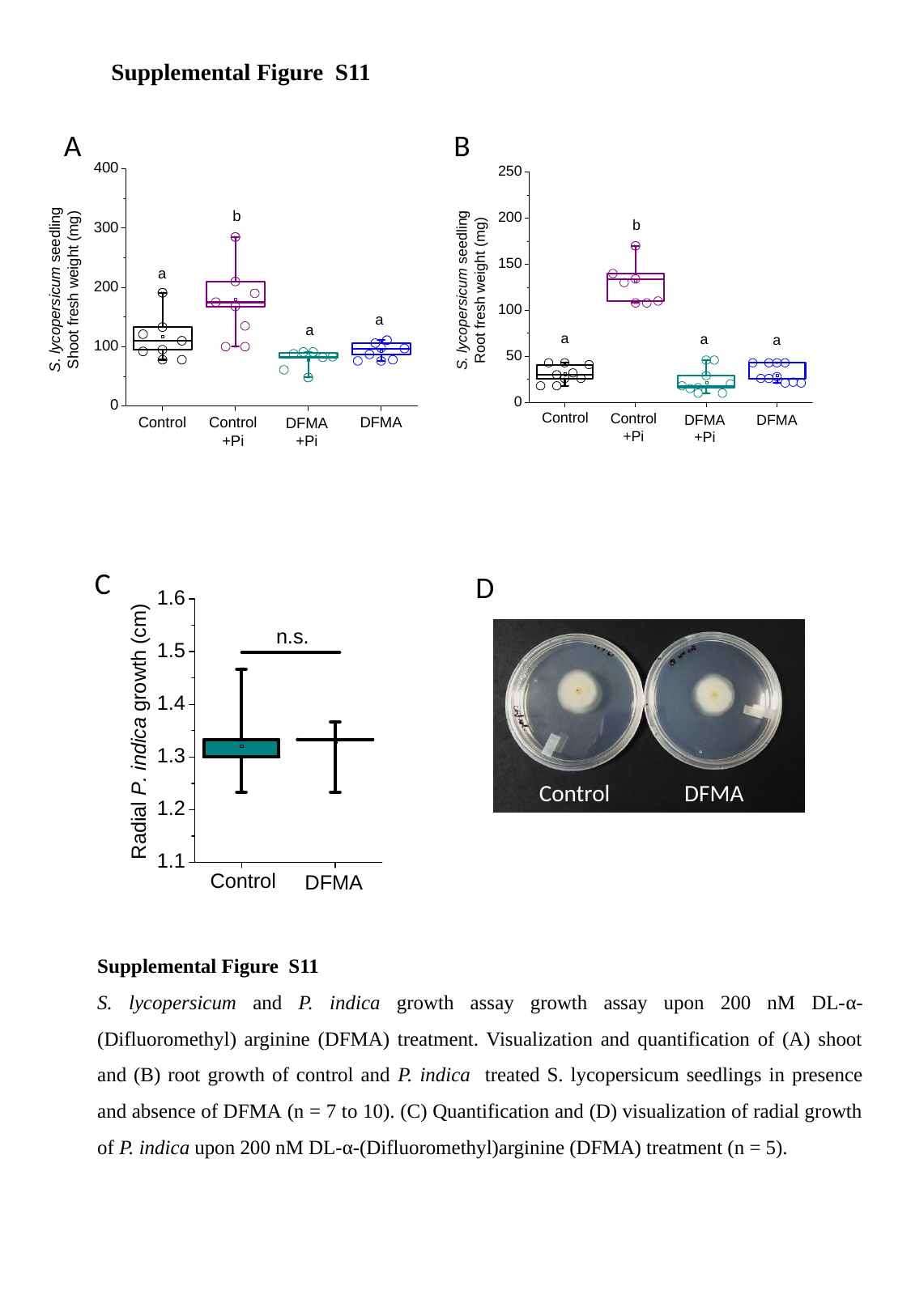

Supplemental Figure S11
A
B
C
D
Control
DFMA
Supplemental Figure S11
S. lycopersicum and P. indica growth assay growth assay upon 200 nM DL-α-(Difluoromethyl) arginine (DFMA) treatment. Visualization and quantification of (A) shoot and (B) root growth of control and P. indica treated S. lycopersicum seedlings in presence and absence of DFMA (n = 7 to 10). (C) Quantification and (D) visualization of radial growth of P. indica upon 200 nM DL-α-(Difluoromethyl)arginine (DFMA) treatment (n = 5).

### Slide 13
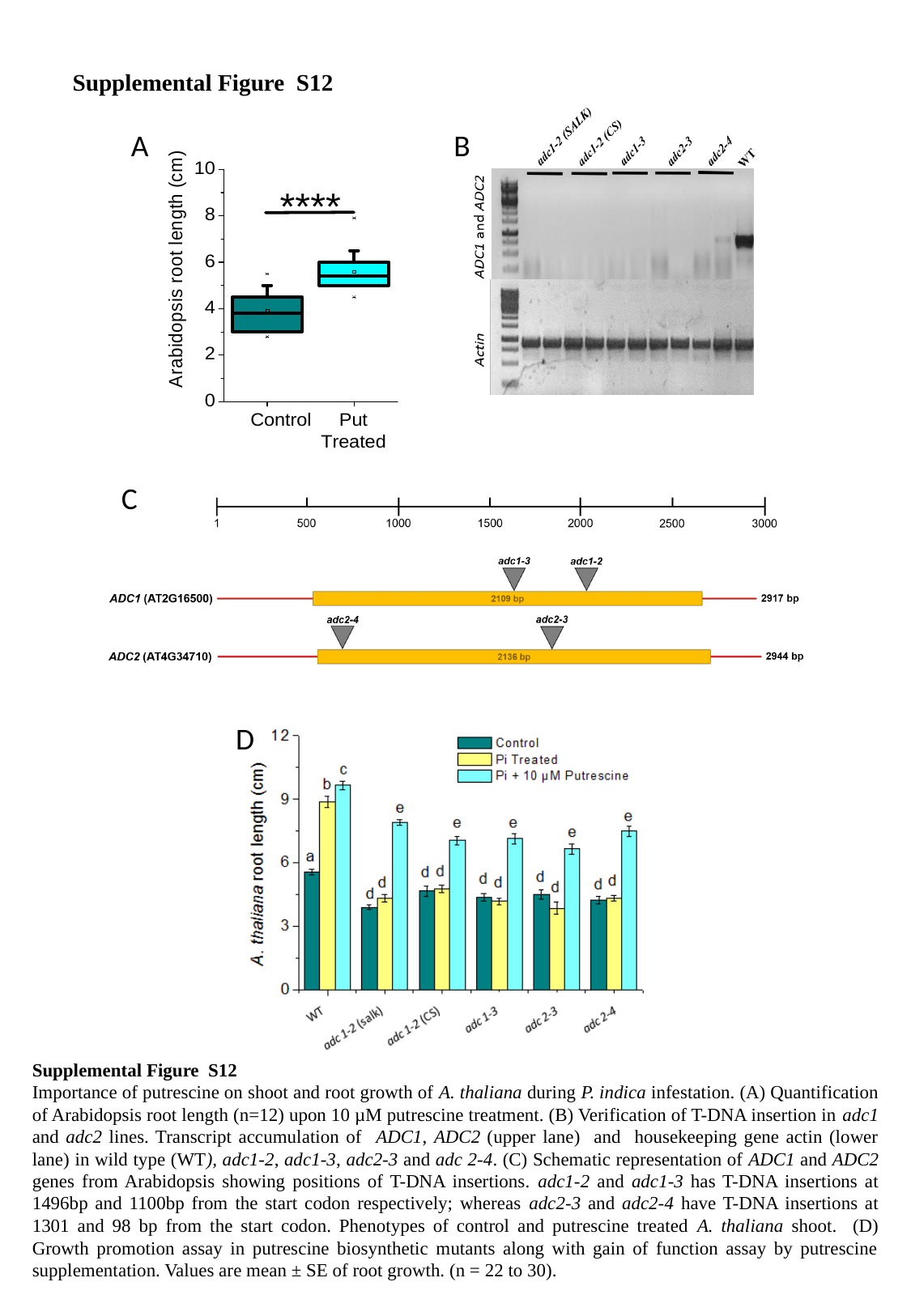

Supplemental Figure S12
B
A
C
D
Supplemental Figure S12
Importance of putrescine on shoot and root growth of A. thaliana during P. indica infestation. (A) Quantification of Arabidopsis root length (n=12) upon 10 µM putrescine treatment. (B) Verification of T-DNA insertion in adc1 and adc2 lines. Transcript accumulation of ADC1, ADC2 (upper lane) and housekeeping gene actin (lower lane) in wild type (WT), adc1-2, adc1-3, adc2-3 and adc 2-4. (C) Schematic representation of ADC1 and ADC2 genes from Arabidopsis showing positions of T-DNA insertions. adc1-2 and adc1-3 has T-DNA insertions at 1496bp and 1100bp from the start codon respectively; whereas adc2-3 and adc2-4 have T-DNA insertions at 1301 and 98 bp from the start codon. Phenotypes of control and putrescine treated A. thaliana shoot. (D) Growth promotion assay in putrescine biosynthetic mutants along with gain of function assay by putrescine supplementation. Values are mean ± SE of root growth. (n = 22 to 30).
